# Microglia undergo transcriptional, translational and functional adaptations to dark and light phases in laboratory mice

**DOI:** 10.1101/2023.11.17.567571

**Authors:** Daniele Mattei, Andranik Ivanov, Jacqueline Hammer, Bilge Ugursu, Sina Schalbetter, Juliet Richetto, Ulrike Weber-Stadelbauer, Flavia Mueller, Joseph Scarborough, Susanne A Wolf, Helmut Kettenmann, Bernd Wollscheid, Dieter Beule, Urs Meyer

## Abstract

Microglia cells are increasingly recognized to contribute to brain health and disease. Preclinical studies using laboratory rodents are essential to advance our understanding of the physiological and pathophysiological functions of these cells in the central nervous system. Rodents are nocturnal animals, and they are mostly maintained in a defined light–dark cycle within animal facilities, with many laboratories investigating microglial molecular and functional profiles during the animals’ light (sleep) phase. However, only a few studies have considered possible differences in microglial functions between the active and sleep phases. Based on initial evidence suggesting that microglial intrinsic clock genes can affect their phenotype, we sought to investigate differences in transcriptional, proteotype and functional profiles of microglia between light (sleep) and dark (active) phases, and how these changes are affected in pathological models. We found marked transcriptional and proteotype differences between microglia harvested during the light or dark phase. Amongst others, these differences related to genes and proteins associated with immune responses, motility, and phagocytosis, which were reflected by functional alterations in microglial synaptic pruning and response to bacterial stimuli. Possibly accounting for such circadian changes, we found RNA and protein regulation in SWI/SNF and NuRD chromatin remodeling complexes between light and dark phases. Importantly, we show that microglial circadian transcriptional changes are impaired in a model of immune-mediated neurodevelopmental disorders. Our findings emphasize the importance of considering circadian factors in studying microglial cells and indicate that implementing a circadian perspective is pivotal for advancing our understanding of their physiological and pathophysiological roles in brain health and disease. This may also open novel avenues towards therapeutic strategies for modulating microglial functions during specific windows of the active or sleep phase.

## 1. Introduction

Microglia cells constitute the immune competent cells of the central nervous system (CNS) (Borst et al., 2021). Their physiological role in the brain extends from pure immunological defense to chaperons of brain circuit formation in development and maintenance of CNS functions in adulthood (Borst et al., 2021). The latter aspect has received increasing attention in recent years, as microglia cells have been shown to participate in a vast array of physiological brain functions in favor of proper circuit stability (Borst et al., 2021; Umpierre and Wu, 2021). Microglia constantly move their processes, and they can sense their environment via an asset of cell-surface receptors termed the sensome (Hickman et al., 2013; Li et al., 2012; Milior et al., 2020). They can sense and respond to neuronal activity and promote synaptic plasticity in an intricate neuro-immune crosstalk (Ferro et al., 2021; Marinelli et al., 2019; Umpierre and Wu, 2021).

Preclinical studies using laboratory rodents are essential to advance our understanding of the physiological and pathophysiological functions of microglial cells in brain health and disease. Rodents are nocturnal animals, and they are mostly maintained under in a defined light–dark cycle within animal facilities, with many laboratories performing analyses of microglial functions during the animals’ light (sleep) phase. There is, however, initial evidence suggesting that microglial functions differ between active and sleep phases (Deurveilher et al., 2021). For example, while initial studies found that pro-inflammatory cytokine expression in rat hippocampi followed a circadian rhythm (Fonken et al., 2015), more recent studies have shown that the expression of the clock genes *Bmal1* (*Arntl*) and *Rev-erbα (Nr1d1)* influence the microglial immune response and phagocytic functions (Griffin et al., 2020, 2019; Wang et al., 2020). However, these studies only looked at selected microglial genes (Fonken et al., 2015), or they analyzed the response to the bacterial endotoxin, lipopolysaccharide (LPS), via mRNA-sequencing. of whole hippocampal tissue(Griffin et al., 2019), but not in a cell-specific manner. Other studies revealed that microglial morphology and process motility also changes between the light and dark phases in response to circadian fluctuations of noradrenaline levels, (Liu et al., 2019; Stowell et al., 2019). For example, Stowell *et al*. observed that noradrenaline-mediated modulation of microglial process motility during the dark (active) phase is associated with a decrease in synaptic plasticity in the visual cortex of of laboratory rodents (Stowell et al., 2019). In line with these findings, Choudhuri *et al*. demonstrated that microglia-mediated phagocytosis of synaptic material is higher during the light (sleep) phase in the prefrontal cortex of adult rats, while Griffin *et al*. found increased microglial synaptic pruning during dark (active) phase in the mouse hippocampus (Choudhury et al., 2020; Griffin et al., 2020). More recently, Corsi *et al*. demonstrated that hippocampal microglia alter their fractalkine receptor (*Cx3cr1*) expression between active and sleep (Corsi et al., 2022), which suggests that circadian factors may play a role in modulating microglial activity through neuron-derived signals.

Taken together, the existing evidence suggests that the functional properties of microglia differ between the active and sleep phases as a result of the influence of clock genes and circadian rhythms. However, a systematic investigation of the transcriptional, proteotype and functional profiles of microglia harvested during the active or sleep phase is still lacking. A multi-modal comparison of microglial profiles during active and sleep phases would help explaining within- and between-study variability in microglial readouts, which may arise due to differences in sampling hours in correlation with facilities’ light-on/off schedules. Moreover, considering circadian factors in microglial functions may also open novel avenues towards their therapeutic modulation, as it may allow more specific chrono-pharmacological targeting of the central immune system. For these reasons, the present study was designed to evaluate the transcriptional, translational, and functional adaptations of microglia to the active (dark) and sleep (light) phases in laboratory mice.

To this end, we focused on hippocampal microglial cells, mainly because the hippocampus is a highly plastic brain region associated with learning and memory processes that are supported by microglial functions (Cornell et al., 2022).

## 2. Results

### 2.1. Microglia undergo transcriptional adaptations to active and sleep phases

#### 3.1.1 Changes in microglial specific genes between the active and aleep phases are accompanied by altered levels of fractalkine and Interleukins

First, we compared the expression of selected microglial genes in freshly isolated hippocampal microglia along with hippocampal cytokine and fractalkine protein levels in samples collected at 1 p.m. (i.e., 4 h after lights were off; = active phase) and 1 a.m. (4 h after lights were on; = sleep phase) (**Fig. 1A**). We found that *P2ry6* and *P2ry12*, both of which are important for purine-mediated microglial phagocytosis and process motility (Bernier et al., 2019; Koizumi et al., 2007), were significantly increased during the active as compared to the sleep phase (**Fig. 1B**). On the other hand, *Siglech*, a microglial gene that is crucial for modulating microglial activation (Konishi et al., 2017), and *Spi1*, a key transcription factor for the microglial myeloid identity, were increased during the sleep phase (**Fig. 1B**). Likewise, we found that hippocampal fractalkine (CX3CL1) and IL-4 were significantly higher during the active as compared to the sleep phase, whereas hippocampal IL-6 levels were lower during the active as compared to the sleep phase (**Fig. 1C**). Hippocampal levels of IL-1β, IL-10 and TNF-α did not change as a function of the dark-light cycle. Taken together, these results provided a first line of evidence suggesting that basal immune microenvironments and microglia-related transcripts in the hippocampus of laboratory mice differ between light and dark phases.

**Figure 1:**
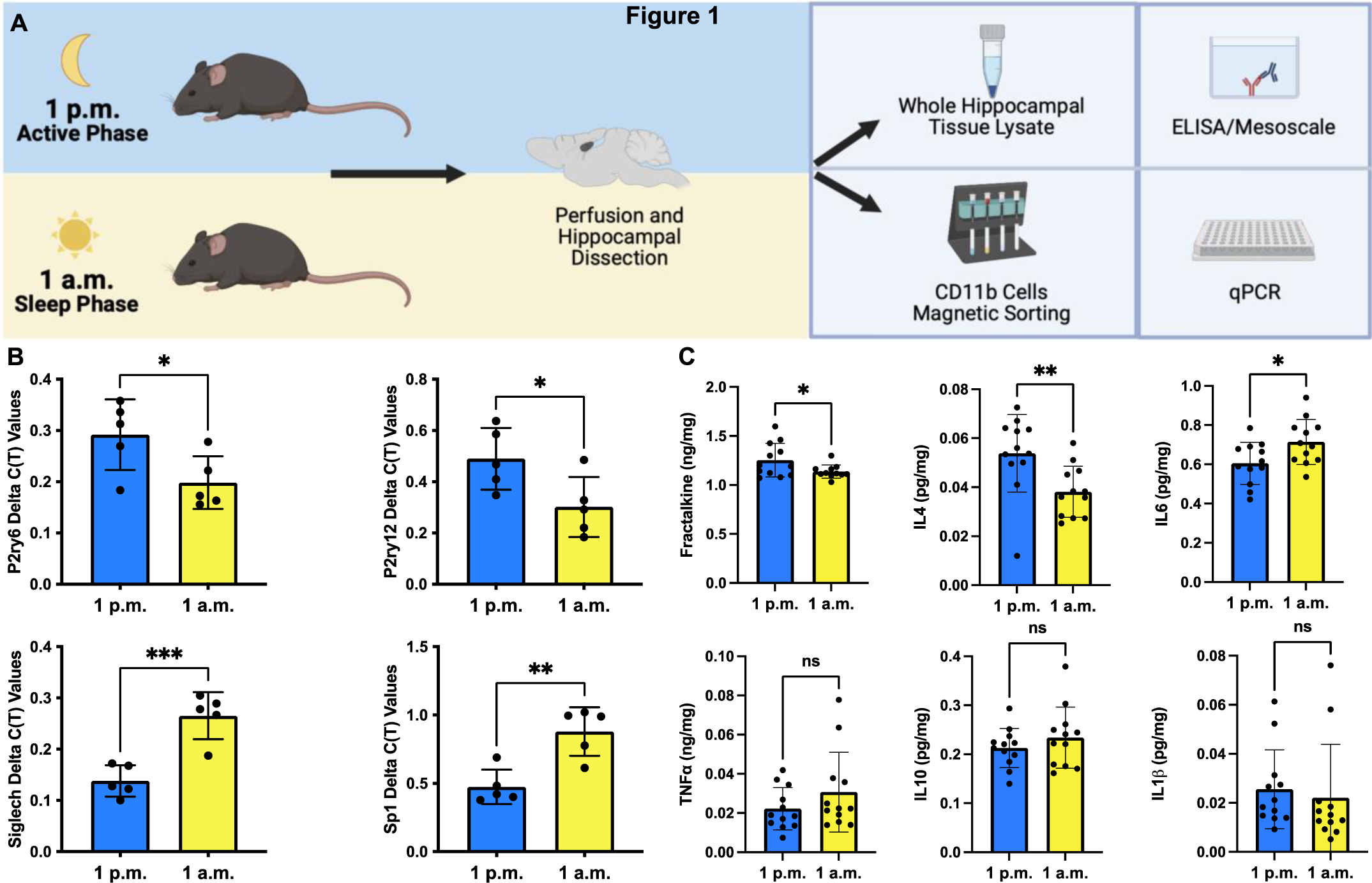
Changes in microglial specific genes between the active and sleep phases are accompanied by altered levels of fractalkine and Interleukins. **A** Schematic representation of the experimental design. Whole hippocampi were retrieved from perfused adult male mice 4h post lights-off (1 p.m., active phase) or 4h post lights-on, (1 a.m., sleep phase) for either microglial isolation or hippocampal protein extraction. Enriched microglial fractions were used for qPCR analysis of specific microglial genes, while hippocampal lysates were used for ELISA and mesoscale measurements of fractalkine and selected cytokines respectively. **B** Gene expression comparison for the microglial genes *P2ry6* (t(8) = 2.44, *p* = 0.041), *P2ry12* (t(8) = 2.5, *p* = 0.037), *Siglech* (t(8) = 5.16, *p* = 0.0009) and *Spi1*(t(8) = 4.15, *p* = 0.0032) between 1 a.m. and 1 p.m. (N = 5 mice/group) **C** Comparison of whole hippocampal protein levels for the chemokine fractalkine (t(20) = 2.4, *p* = 0.026, N = 11 mice/group) and cytokines interleukin-4 (IL-4) (t(22) = 2.87, *p* = 0.0089), interleukin-6 (IL-6) (t(22) = 2.4, *p* = 0.025), tumor necrosis factor-α (TNF-α), interleukin-10 (IL-10) and interleukin-1β (IL-1β) between 1 a.m. and 1 p.m. (N = 12 mice/group). Error bars represent the mean ± standard deviation. Unpaired two-tailed t-test **\****p*< 0.05, \*\**p*< 0.01, \*\*\**p*< 0.001. Images created with Biorender.com.

#### 3.1.2 Microglia cells undergo transcriptional adaptations between the active and sleep phases

To further substantiate this notion, we performed unbiased, genome-wide RNA sequencing using hippocampal microglia that were collected at 1 p.m. (i.e., 4 h after lights were off; = active phase) and 1 a.m. (4 h after lights were on; = sleep phase) (**Fig 2A**). These analyses revealed marked differences between the transcriptomic profiles of microglia collected during the active or sleep phase. Indeed, compared to microglia collected during the active phase, 1328 and 231 genes were up- and downregulated, respectively, in microglia collected during the sleep (light) phase as compared to the active (dark) phase (**Fig. 2B**, **Table S1**).

**Figure 2:**
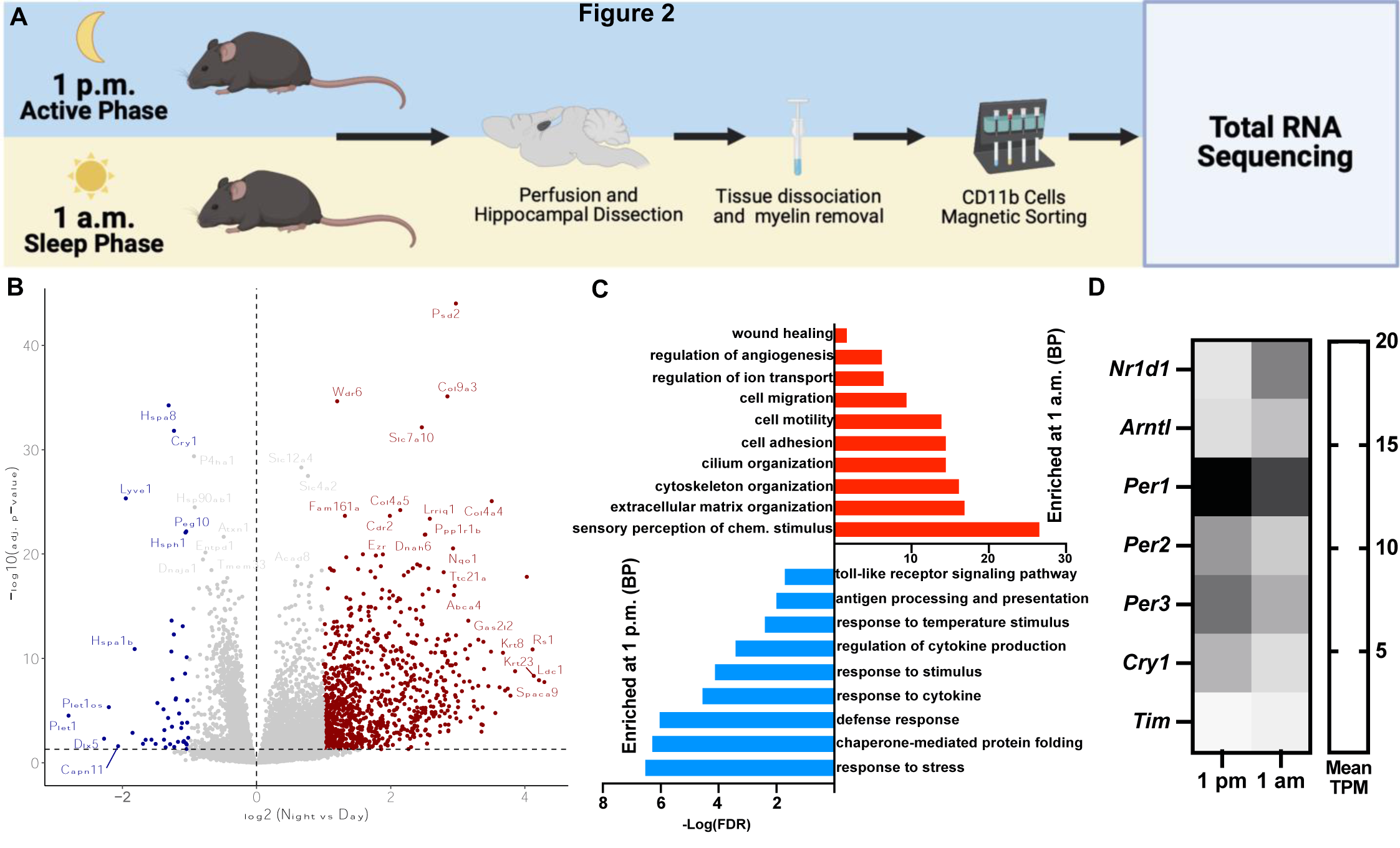
Microglia cells undergo transcriptional adaptations between the active and sleep phases. **A** Schematic representation of the experimental design. Whole hippocampi were retrieved from perfused adult male mice at either 1 p.m. (active phase) or 1 a.m. (sleep phase) for CD11b cell enrichment and subsequent total-RNA-sequencing. **B** Volcano plot representing the differential expression in microglial genes between 1 p.m. and 1 a.m. A total of 1559 genes were found to be differentially expressed between 1 a.m. and 1 p.m., (Log2FC cut-off of 0.5, adjusted *p-*value of 0.05, N = 15-16 animals/group. **C** -Log(FDR) values for selected biological processes from the gene ontology analysis of the up- and downregulated genes between 1 a.m. and 1 p.m. **D** Heatmap displaying changes in clock gene expression in transcripts per million (TPM) between 1 a.m. and 1 p.m. Images created with Biorender.com.

Besides others, genes that are crucial for phagocytotic activity (Sierra et al., 2013), including *Dock*-family genes, *Mertk, Rhob, Elmo1, Syk, Hck, Csk* and *Rac3* (**Fig. 3B**, **Fig. S4F**), were found to be upregulated during the active phase, suggesting that the transcriptional correlates of phagocytotic activity follow a circadian regulation. Consistent with the findings of increased hippocampal fractalkine (CX3CL1) levels during the active phase (Fig. S1C), we also found the active phase to be associated with increased expression of *Adam10* (**Fig. S1C**), which encodes the enzyme responsible for fractalkine cleavage (Gunner et al., 2019). Other DEGs that were upregulated during the active phase were genes related to chaperone-mediated protein folding (**Fig. S4C-D)**, innate immune response (**Table S2**), antigen presentation, Toll-like receptor signaling pathways (**Fig.S4E**), and phagocytic receptors (**Fig. 2C**, **Table S1**). The latter included *Fcgr1*, *Fcgr*3 *C5ar1* and C*5ar2* found enriched during the active phase (**Fig S4E-F**), along with genes characterizing disease associated microglia (DAM) signatures observed in β- amyloid plaque-associated microglia (**Table S2**), (Grubman et al., 2021). On the other hand, specific DEGs that were upregulated during the sleep (light) phase were annotated with cell adhesion (**Fig. S4A-B**), motility, angiogenesis, and extracellular matrix (ECM) remodeling (**Fig. 2C**, **Table S1**). The latter involved e.g., *Adamts4, Mmp14,* and *Ctsc* (**Fig. S1**), relevant for microglial ECM-remodeling and engulfment in favor of synaptic plasticity (Nguyen et al., 2020).

**Figure 3:**
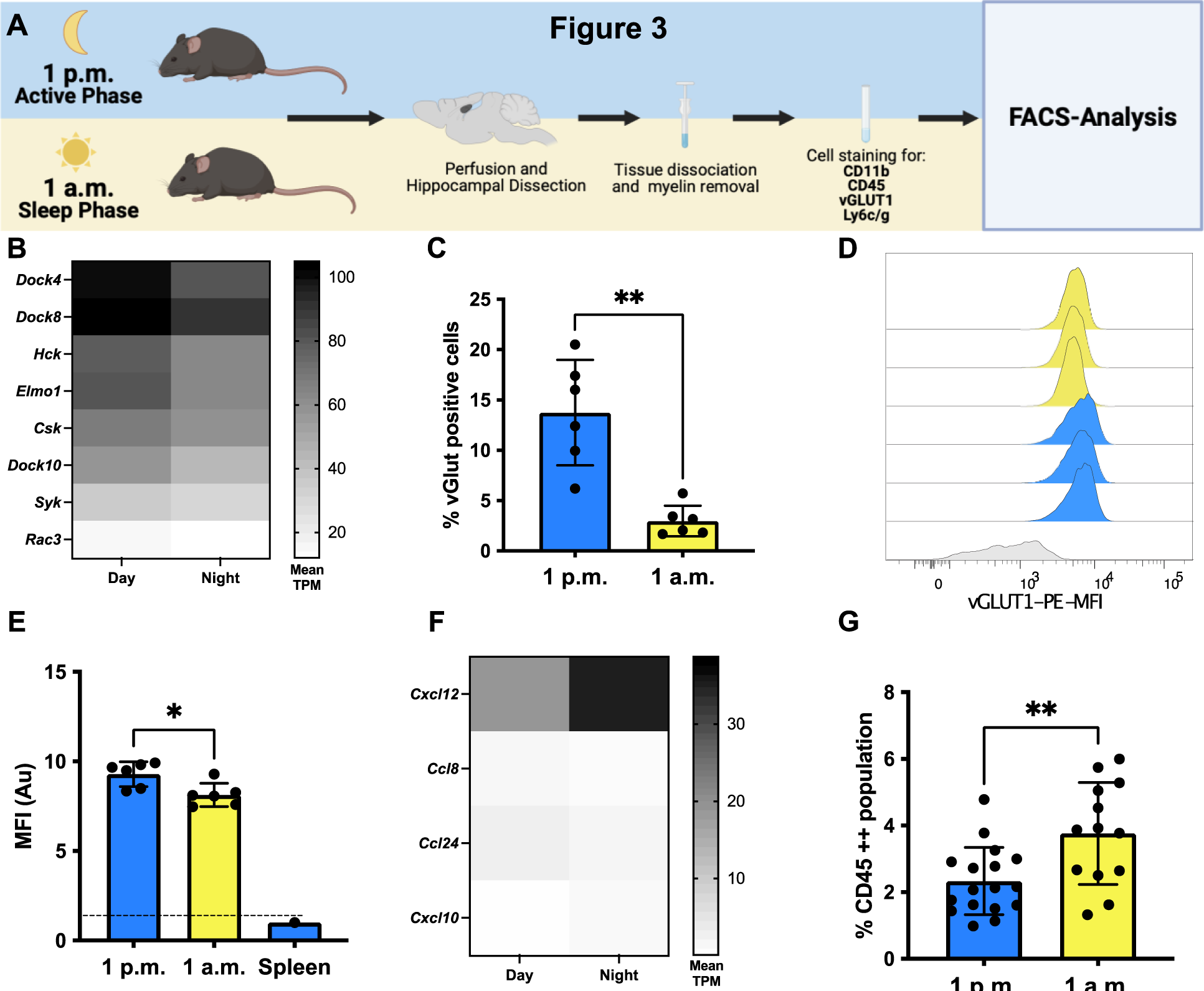
Microglia display increased synaptic pruning during the active phase In the adult hippocampus. **A** Schematic representation of the experimental setup. Whole hippocampi were retrieved from perfused adult male mice either 4h post lights-off (active phase) or 4h post lights-on (sleep phase) for CD11b cell enrichment and subsequent flow cytometry analysis of vGlut1-posive cells. **B** Heatmap showing the differential expression (in average transcripts per million, TPM), for selected genes associated with microglial phagocytic functions from the total-RNA-seq of microglia 4h post lights-off or 4h post lights-on (Fig. 2, 1 a.m. versus 1 p.m., Only genes that were significantly differentially expressed are represented, adj. *p*-value < 0.05) **C** Scatter plot showing the difference in percentage of vGlut1-positive microglia cells between the active and sleep phase, 4h post lights-off and on respectively, (unpaired t-test, t(5.84) = 4.84, p = 0.0031, N = 6/group). **D** Histogram showing the microglial vGlut1-PE mean fluorescence intensity normalized to spleen cells as negative control (lower panel). **E** Scatter plot showing the area under the curve (Au) for the microglial vGlut1 mean fluorescence intensity during the active (4h after light-off) and sleep (4h post light-on) phases, normalized to spleen cells as negative controls (unpaired t-test, t(10) = 2.98, p = 0.014, N = 6/group). **F** Heatmap showing differential expression (in average transcripts per million, TPM), for selected genes coding for chemokines (from the total-RNA-seq of microglia 1 p.m. versus 1 a.m.). Only genes that were significantly differentially expressed are represented (adj. *p*-value < 0.05). **G** Scatter plot comparing the percentage of CD45^hi^ cell population in the hippocampi of adult mice 4h after light-off (N = 17) and 4h post light-on (N = 13, unpaired t-test, t(28) = 3.08, *p* = 0.0046). Error bars represent the mean ± standard deviation. **\****p*< 0.05, \*\**p*< 0.01. Images created with Biorender.com.

Confirming our initial qRT-PCR analysis (**Fig. 1B**), we also found lower transcript levels of *P2ry12* and *P2ry6* in hippocampal microglia collected during the sleep phase (**Fig. S1A**). These changes were accompanied by decreased expression of *Entpd1* (**Fig. S2A**), which encodes for CD39, a membrane bound enzyme responsible for converting extracellular ATP to ADP. The latter stimulates P2Y12 receptors and promotes microglial process motility (Illes et al., 2020). Furthermore, the transcripts of various neurotransmitter receptors, including serotonergic, cholinergic, and adrenergic receptors (**Fig. S3**), as well as transcripts of several heat-shock proteins (**Fig. S4C-D**), were significantly altered between active and sleep phases. Notably, several clock genes were also found to be expressed differently in hippocampal microglia that were collected during the sleep as compared to the night phase (**Fig. 2D**), confirming previous reports of circadian effects on clock gene expression in microglia (Griffin et al., 2020, 2019; Wang et al., 2020). Taken together, these results demonstrate that hippocampal microglia undergo widespread transcriptional adaptations between light and dark phases in laboratory mice.

### 3.2. Microglia display increased excitatory synaptic pruning during the dark phase

Fractalkine is a “find me” signal that can lead to microglial phagocytosis of synaptic material, (Paolicelli et al., 2014). We found increased levels of fractalkine during the active phase (**Fig. 1C**) accompanied with an enrichment in phagocytosis-associated genes (**Fig. 3B**) (Sierra et al., 2013). Altogether, these data suggest a higher level of synaptic pruning during the active phase. We therefore assayed whether the present microglial transcriptional profile and increased fractalkine levels were followed by an increased synaptic engulfment during the active phase. For the purpose, we employed a recently published protocol to detect synaptic material within microglia cells via fluorescent associated cell sorting (FACS) analysis, (Brioschi et al., 2020). We collected microglia from adult mouse hippocampi 4h post light-on and 4h post lights-off as displayed in **Fig. 3A**. We observed a significant increase in percentage of microglia positive for the excitatory presynaptic marker vesicular glutamate transporter-1 (vGlut1) during the active phase (**Fig. 3C**), accompanied by a significantly higher vGlut1 mean fluorescence intensity (MFI, **Fig. 3D-E**, the complete gating strategy is displayed in **Fig. S5**). This is in accordance with Griffin et al. that found a higher number of synaptophysin-positive microglial inclusions during the active phase in the adult mouse hippocampus (Griffin et al., 2020). For this experiment, we distinguished microglia cells from peripheral monocytes by the high expression of CD11b and low expression of CD45. We have previously shown that a limited number of peripheral monocytes are allowed into the brain parenchyma in support of adult hippocampal neurogenesis (Möhle et al., 2016). The RNA-seq also revealed an increased expression of chemokines *Cxcl12* and *Cxcl10* during the sleep phase, accompanied by a decreased expression in *Ccl8* and *Ccl24*, (**Fig. 3F**), which may reflect on changes in chemoattraction of peripheral monocytes. We therefore investigated whether monocytes identified as CD45^hi^ and CD11b^low^ were more abundant during either phase, (**Fig. 3G**). Of note, animals were perfused to remove peripheral immune cells from the brain vasculature. Interestingly, we found an increased proportion of CD45^hi^ and CD11b^low^ during the sleep phase (**Fig. 3G**).

### 3.3. The microglial response to bacterial lipopolysaccharide changes in intensity between the active and sleep phases

#### 3.3.1 The microglial transcriptional response to lipopolysaccharide changes in intensity between the active and sleep phases

Previous studies indicate a higher peripheral innate immune response to LPS during the sleep phase in rodents (Keller et al., 2009). As we observed a significant phase-dependent microglial regulation TLR-related signaling pathway and in *Tlr4* expression between the active and sleep phases (**Fig. 2C**, **Fig. S4E**), we sought to study whether the *in vivo* microglial response to the TLR4 agonist LPS changes between the light and dark phase. LPS was administered intraperitoneally (1mg/Kg) to adult male mice either two hours post light-on (11.00 a.m.) or post light-off time (11.00 p.m.). Hippocampal microglia cells were collected four hours post injection (03.00 p.m. and 03.00 a.m. respectively, **Fig. 4A**). Of note, to avoid technical and batch effects and to be able to best compare the LPS responses, the injections were performed within the same 24h, with the same lot of mice and LPS batch. Furthermore, RNA isolation and library preparation were performed simultaneously for all groups.

**Figure 4:**
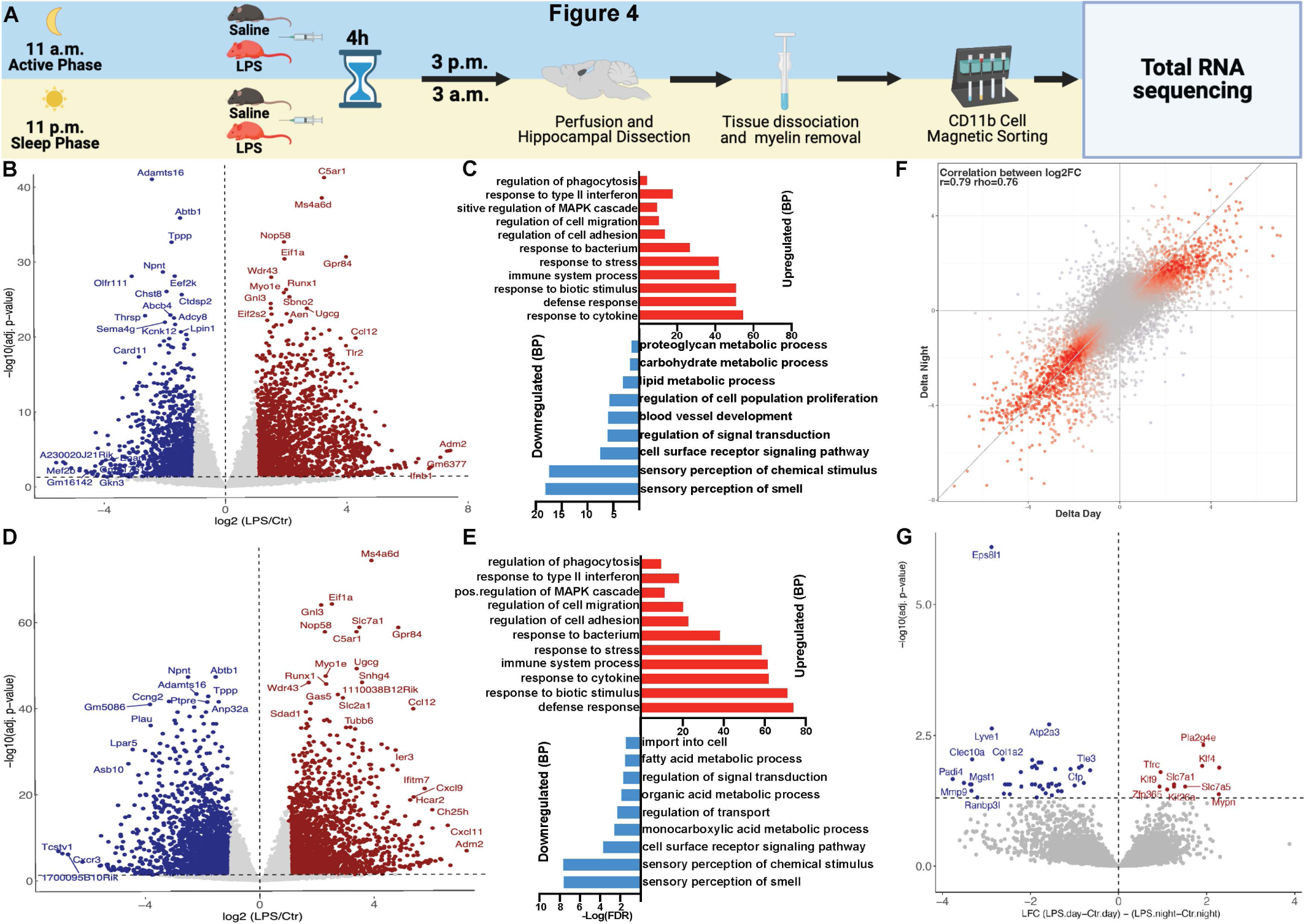
The microglial transcriptional response to lipopolysaccharide changes in intensity between the active and sleep phases. **A** Schematic representation of the experimental setup. Adult male mice were injected intraperitoneally with either lipopolysaccharide (LPS) or saline solution at 11 a.m. (2h post lights- off, active phase) or 11 p.m. (2h post lights-on, sleep phase). Hippocampi were retrieved 4h post injection for microglial isolation and subsequent total-RNA-sequencing, to compare the microglial immune response to LPS during the active and sleep phase. **B** Volcano plot displaying the differential expression between microglia from animals injected with LPS versus saline during the active phase (3 p.m., N = 3 mice/group). A total of 3286 genes were found to be differentially expressed in response to LPS administered during the day (Log2FC cut-off of 1, adj. *p*-value < 0.05). **C** -log(FDR) values for selected biological processes associated with the up- and downregulated genes in response to LPS during the active phase. **D** Volcano plot displaying the differential expression between microglia from animals injected with LPS versus saline during the sleep phase (3a.m., N = 2-3 mice/group). A total of 2799 genes were found to be differentially expressed in response to LPS administered during night (Log2FC cut-off of 1, adj. *p*-value < 0.05). **E** -log(FDR) values for selected biological processes associated with the up- and downregulated genes in response to LPS during the night. **F** Disco plot for the correlation between the absolute Log2-fold changes in gene expression in response to LPS during the active (x-axis) and sleep (y-axis) phase. **G** Volcano plot showing genes significantly regulated as an effect of time of LPS injection (Adj. *p*< 0.05). Images created with Biorender.com.

Total RNA-sequencing revealed that peripheral LPS during the active phase induced a deregulation of 3286 protein coding genes (**Fig. 4B-C**, 1675 up- and 1611 downregulated genes). LPS administration during the sleep phase caused a deregulation of 2799 protein coding genes (**Fig. 4D-E**, 1674 up- and 1125 downregulated genes, **Table S3** contains the complete differential expression and GO-analysis). GO-analysis revealed similar pathways to be deregulated in both time-points (**Fig. 4C, E**), however, the LPS response during the sleep phase showed a stronger enrichment for immune-related pathways such as defense response, response to cytokine, and response to bacterium, as compared to the LPS response during the active phase (**Fig. 4C, E**), despite the similar number of upregulated genes in both phases. This enrichment was observable also at different Log2FC cut-offs (**Table S3**). This suggests a higher innate immune reactivity during the sleep phase, as observed in peripheral macrophages in response to LPS (Keller et al., 2009) and similar to what was found in whole hippocampal tissue in response to systemic LPS (Fonken et al., 2015). To better understand the effect of time on the LPS response we computed directly the contrast in effects by subtracting the observed change in the LPS group during the day (3 p.m.) from the control group during the day (3 p.m.), and similarly, by subtracting the observed change in the LPS group during the night (3 a.m.) from the control group during the night (3 a.m.), (Delta = (LPS_3p.m. vs. Control_3p.m.)- (LPS_3a.m. vs. Control_3a.m.)). Despite overall high similarity in expression changes in either phase (**Fig. 3D**), we found 56 genes to be specifically regulated by time of injection, (**Fig. 4F-G**, **Table S3**). The latter genes were associated with processes such as regulation of leukocyte migration, extracellular matrix remodeling and immune activation (**Fig. S6A-B**, **Table S3**).

#### 3.3.2 The microglial proteotype response to lipopolysaccharide changes in intensity between the active and sleep Phases

We thus generated a new set of animals to study whether hippocampal microglia cells follow a similar pattern at the translational level (**Fig. 5A**) via a Liquid Chromatography– Tandem Mass Spectrometry (LC–MS/MS) Analysis. LPS administered at 11 a.m. (active phase) induced the deregulation of 875 microglial proteins, (**Fig. 5B-C**, 372 up- and 503 downregulated). LPS injection at 11 p.m. (sleep phase) induced the deregulation of 385 proteins, (**Fig. 5D-E**, 106 up- and 279 downregulated, **Table S3** contains the complete proteotype and GO-analysis). GO-analysis analysis revealed that 4h post LPS injection during the sleep phase there was a greater enrichment in proteins associated with an immune response as compared to the active phase (**Fig. 5C, E**), corroborating our findings at the RNA level. Enhanced enrichment in immune-related proteins in response to LPS during the sleep phase was observed irrespective of Log2FC cut-off (**Table S3**). LPS injection during the active phase induced a stronger enrichment in proteins presumably associated with the mounting of an immune response, such as translation and RNA metabolic processes as well as chromatin remodeling and epigenetic changes (**Fig. 5C, E**, **Table S3**). Since proteins are the final effectors of a given transcriptional response, this data suggests that microglia cells react faster to LPS during the sleep phase. Besides regulation in classical immune and metabolic pathways, the response to LPS induced a downregulation in olfactory receptors genes such as *Olfr110* and *Olfr111* resulting in enrichment of BPs such as sensory perception of smell (**Fig. 4B-E**), that was corresponded at the proteotype level (**Fig. 5C, E**). Expression of olfactory receptors by innate immune cells such as microglia and macrophages have been only recently reported. Such receptors regulate immune responses, especially to bacterial stimuli, (Lee et al., 2020; Orecchioni et al., 2022). Of note, as for the transcriptional evaluation, injections for this experiment were performed within the same 24h-period, with the same LPS batch and lot of animals, to minimize batch and technical biases. Moreover, the proteins were extracted in parallel for all groups.

**Figure 5:**
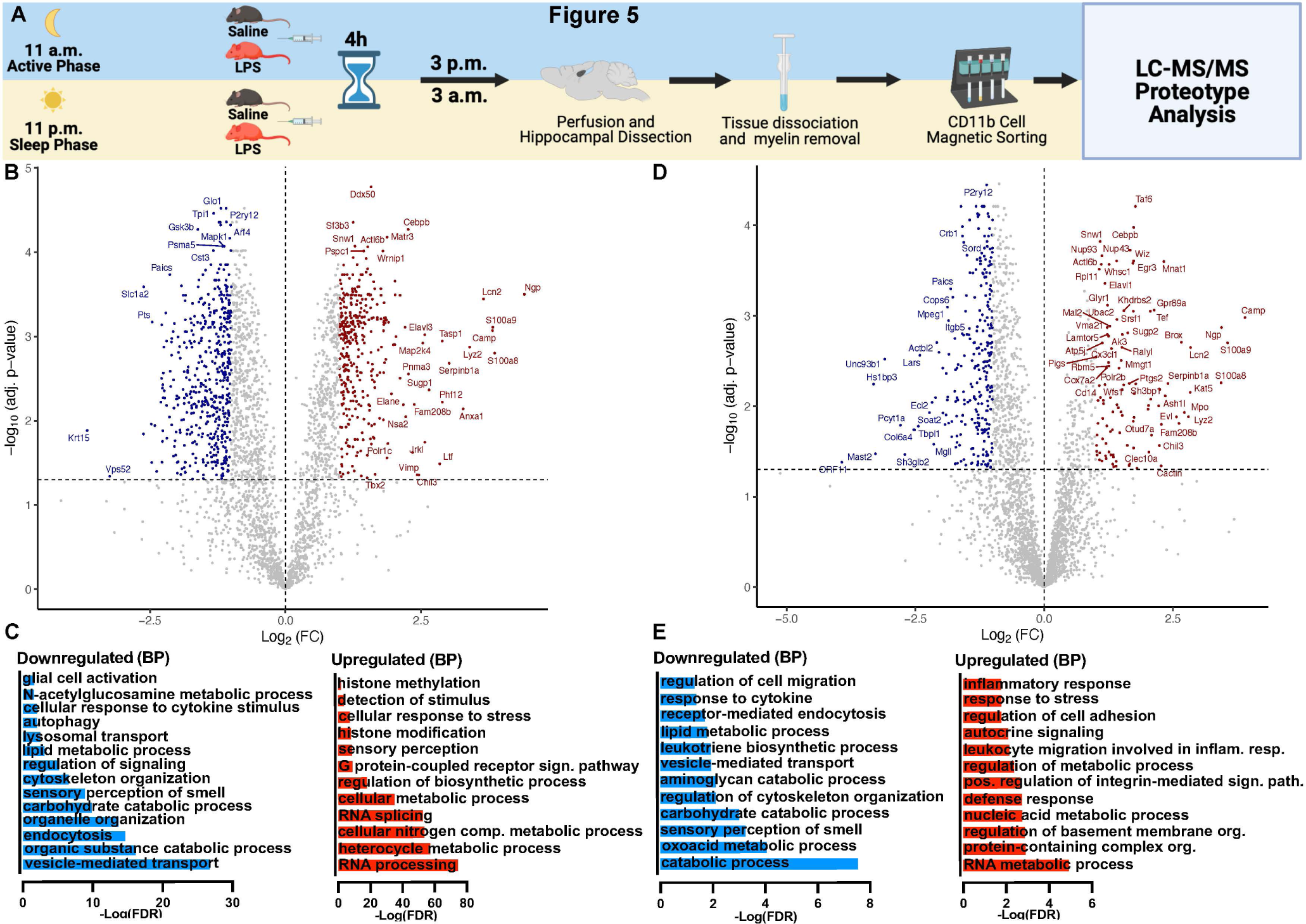
The microglial proteotype response to lipopolysaccharide changes in intensity between the active and sleep Phases. **A** Schematic representation of the experimental setup. Adult male mice were injected intraperitoneally with either lipopolysaccharide (LPS) or saline solution at 11 a.m. (2h post lights- off, active phase) or 11 p.m. (2h post lights-on, sleep phase). Hippocampi were retrieved 4h post injection for microglial isolation and subsequent Liquid Chromatography–Tandem Mass Spectrometry (LC-MS/MS) analysis to compare the microglial immune response to LPS during the active and sleep phase. **B** Volcano plot showing the differentially expressed proteins in response to LPS administered during the active phase (3 p.m., N = 4 mice/group). 875 proteins were differentially expressed 4h post LPS injection (Log2FC cut-off of 1 and adj. *p-*value < 0.05). **C** -log(FDR) values for selected biological processes associated with the up- and down-regulated proteins in response to LPS during the active phase. **D** Volcano plot showing the differentially expressed proteins in response to LPS administered during the sleep phase (3 a.m., N = 4 mice/group). 385 proteins were differentially expressed 4h post LPS injection during the sleep phase (Log2FC cut-off of 1 and adj. *p-*value < 0.05). **E** -log(FDR) values for selected biological processes associated with the upregulated proteins in response to LPS during the sleep phase. Images created with Biorender.com.

### 3.4. The microglial transcriptional and proteotype regulation between the active (3p.m.) and sleep (3 a.m.) phases

To further investigate microglial transcriptional and proteotype differences between the active and sleep phases, we utilized the data from control animals generated to study the LPS response (**Fig. 3A**) to compare the transcriptional and proteotype profile of microglia cells between 3 p.m. and 3 a.m., hence 6h post light-off and -on respectively. We found 182 protein-coding DEGs between these two timepoints (56 upregulated and 126 downregulated**, Fig S7A**, **Table S4** contains the complete differential expression and GO-analysis). Similar to the changes found between 1 a.m. and 1 p.m., microglia showed transcriptional regulation in BPs associated with cell migration, motility, adhesion, and angiogenesis (**Fig. S7B**). Additionally, genes associated with neuronal processes and myelination were enriched at 3 p.m. (**Fig. S7B**). Microglia have been shown to express neuronal and oligodendrocyte genes, albeit to a lower extent, and the function of such transcripts in microglia remains elusive, (Grassivaro et al., 2020; Solga et al., 2015). Notably, genes associated with cell differentiation were enriched during the sleep phase (**Fig. S7B**). Proteotype analysis of microglia freshly isolated from control mice at 3 p.m. and 3 a.m. revealed 481 proteins differentially expressed (251 up- and 230 downregulated proteins, **Fig S7C**, **Table S4** contains the complete proteotype analysis). We found proteins associated with cell-adhesion and cell differentiation to be enriched at 3 a.m., in correspondence with the transcriptional analysis (**Fig. S7B**, **D**), while cell motility- and angiogenesis-related proteins were enriched at 3 p.m. Also, corresponding the RNA-analysis, neuronal-related processes such as cell projection organization, were enriched at 3p.m. (**Fig. S7D**). Looking deeper into hippocampal microglial changes in protein expression we found increased levels of C1QA and C1QC in the active phase, confirming what we found transcriptionally at 1 p.m., as reported in **Table S5**. The latter table summarizes the transcriptional and proteotype changes in selected genes and proteins relevant to microglial states and functions. Higher complement component proteins (**Table S5**) along with increased fractalkine during active (**Fig. 1C**) may participate to the higher rate of synaptic pruning observed during the active phase (**Fig. 3**). In line with this, we found that phagocytic marker CD68 and the proteins paxillin and cofilin-2 were also enriched during the active phase, along with CDC42 (**Table S5**). The latter are important drivers of cytoskeletal changes necessary for phagocytosis to occur, (Gitik et al., 2014; López-Colomé et al., 2017). Concomitantly, we found an active phase-associated enrichment in myosin proteins MYH10 and MYH14, required for proper cell motility and phagocytosis, (Porro et al., 2021), along with chaperonin-containing TCP1 subunits 2, 3 and 5 (CCT2, CCT3 and CCT5, **Table S5**). CCT chaperonins are critical for proper myosin and actin folding and fibril formation, (Srikakulam and Winkelmann, 1999). Importantly, CCT5 interacts with F-actin enabling proper MERTK-dependent phagocytic functions (Feng et al., 2022). Furthermore, CD81 was enriched during the active phase (**Table S5**), a protein relevant for microglia motility and myelin phagocytosis (Martins et al., 2019). The proteotype analysis also revealed a decrease in P2Y12 and CD39 levels during the sleep phase, as were the corresponding transcripts (**Table S5**), confirming a circadian regulation of the microglial purinergic system. Finally, we report circadian regulation of genes and proteins associated with glucose and lipid metabolism along with mitochondrial functions (**Table S5**), indicative of metabolic reorganization between phases. Next, we sought to understand the potential machinery behind the microglial circadian molecular and functional changes. We observed a sleep phase-associated enrichment in BPs such as chromatin remodeling and nucleosome organization (**Fig. S7D**). Accordingly, we found marked protein enrichment for several components of the chromatin remodeling complexes (CRCs) SWItch/Sucrose Non-Fermentable (SWI/SNF) and Nucleosome Remodeling and Deacetylase (NuRD, **Table S5** and **S6**) at 3 a.m. The latter complexes can swiftly change the chromatin architecture thereby mediating changes in cellular state and are emerging as key players of innate immune cells’ ability to flexibly alter their states, (Gatchalian et al., 2020; Ramirez-Carrozzi et al., 2006). CRC activity is guided by epigenetic readers, i.e., proteins that binds to specific histone and DNA epigenetic marks and recruit CRCs to appropriate chromatin sites. We found a sleep phase-associated protein enrichment in epigenetic readers, amongst which CHDs, (part of the NuRD complex) and bromodomain-containing factors (BRDs, recruiters of the SWI/SNF complex), at baseline and in response to LPS (**Table S6**). These are particularly important transcriptional regulators of the response to LPS in macrophages and microglia, (Bao et al., 2017; Ramirez-Carrozzi et al., 2006; Wang et al., 2018). We further found that following LPS injection, protein regulation of CRCs and epigenetic readers, was more pronounced during the active phase (**Table S6**), as also reflected by the associated BPs (**Fig.3I**). This is likely because these proteins are already enriched during the sleep phase (**Table S6**), indicating a higher microglial state of flexibility.

### 3.5. Microglia cells display a deregulated phase-associated transcriptional adaptation in the maternal immune activation model of neurodevelopmental disorders

Microglia cells have garnered extensive attention in the field of neurodevelopmental disorders, including conditions like schizophrenia, (Mattei and Notter, 2020). Adverse events during pregnancy such as maternal immune activation (MIA) represent environmental risk factors for such disorders (Meyer, 2019). This is modeled in rodents via the injection of the viral mimic Poly(I:C) during pregnancy (Meyer, 2019). The latter, a synthetic double-stranded RNA, binds to the Toll-like receptor 3 (TLR3) eliciting a viral-like immune response, (Meyer, 2019). In prior studies, we demonstrated that injection of Poly(I:C) during gestational day 17 (GD17) yields adult offspring presenting several hippocampal abnormalities, (Giovanoli et al., 2015). We have also shown that offspring from dams injected with Poly(I:C) display aberrant transcriptional profiles in hippocampal microglia, (Mattei et al., 2017). Of note, for the latter study, animals were hosted in a light:dark cycle, and microglia were collected during the light phase (Mattei et al., 2017). Thus, we generated GD17 Poly(I:C) adult offspring (Poly animals) to extend the transcriptional analysis to look at 1) the microglial transcriptional differences between the light (sleep) and dark (active) phases in adult Poly animals (within-group comparison: Poly-3p.m. versus Poly-3a.m., 2) to compare the adult Poly microglial transcriptional profile to control animals with respect to the light and dark phase (between-group comparisons: Poly-3p.m. versus Control-3p.m. and Poly-3a.m. versus Control-3a.m.) and 3) to compare the response to LPS between the active and sleep phases in microglia from Poly animals, as displayed in **Fig. 6**. When comparing the microglial transcriptional changes between the sleep and active phase in Poly animals (Poly-3a.m. vs. Poly-3p.m.), we found 2527 DEGs, (1399 upregulated and 1128 downregulated, **Fig 7A**, **Table S7**). As a comparison, for the same microglial contrast in control animals, we found 183 DEGs (control day 3 a.m. versus control night 3 p.m., **Fig. S7**, **Table S4**). The direct contrast of PolyI:C effects on adult microglial transcriptional changes between the active and sleep phases (Delta = (PolyI:C_3p.m. vs. Ctr_3p.m.)-(PolyI:C_3a.m. vs. Ctr_3a.m.)) reveals 1420 significant protein coding genes (574 upregulated and 845 downregulated genes, **Supplementary Fig. S8, Table S7,** analogous to figure **Fig. 3D-E**). These genes subject to the MIA effect were associated with BPs such as immune response, phagocytosis, cell adhesion and neurogenesis (**Supplementary Fig. S8, Table S7).** This suggests that the physiological adaptation between the light and dark phase is profoundly affected in this model of neurodevelopmental disorders. Genes associated with innate immune functions, including cytokine production and phagocytosis were enriched during the active phase (**Fig. 7A**, **Table S7**), deviating from what we found in control animals for the same comparison (**Fig. S7C**, **Table S4**). Genes associated with cell adhesion, which in control conditions were enriched at 3 a.m. (**Fig. S7C**) were instead strongly enriched at 3 p.m. in adult Poly microglia. Genes linked to neurogenesis were enriched at 3 p.m., alike in control conditions (**Fig S7C**) albeit to higher extent (**Fig 7A**, **Table S7**). Notably, genes associated with sensory perception of smell where highly enriched at 3 a.m. in Poly microglia. As mentioned in the result section 3.3, olfactory receptors regulate innate immune responses (Orecchioni et al., 2022), and they were subject to expression changes at both the RNA and protein level in response to LPS (**Fig. 4C**, **E**, **Fig. 5C**, **E**). When comparing the transcriptional profile of microglia collected during the active phase from Poly animals and controls (Poly-3p.m. versus Control-3p.m., **Fig. 7B**), we found 901 DEGs (298 upregulated and 603 downregulated, **Fig. 7B**, **Table S7**). The upregulated genes were related to BPs such as immune response, regulation of IL1 and TNF production and lipid metabolism, whilst the downregulated genes were associated with cell adhesion, extracellular matrix organization as well as regulation of synaptic signaling (**Fig 7B**). When comparing the transcriptional profile of microglia collected during the sleep phase from Poly animals and controls (Poly-3a.m. versus control-3a.m., **Fig. 4D**), we found 250 DEGs (157 upregulated and 93 downregulated). The upregulated genes were associated with BPs such as cell development, glia cell differentiation, cell adhesion and cell development (**Fig. 7C**). The downregulated genes were associated with angiogenesis, cell motility, chemotaxis, and immune system processes (**Fig. 7C**). Importantly, the latter pathways were found deregulated also in our previous transcriptional analysis, (Mattei et al., 2017), where microglia were collected from Ploy and control animals during the sleep phase. We thus show that the same comparison performed with microglia cells collected during the active phase can reveal transcriptional deregulations different from the sleep phase.

**Figure 6:**
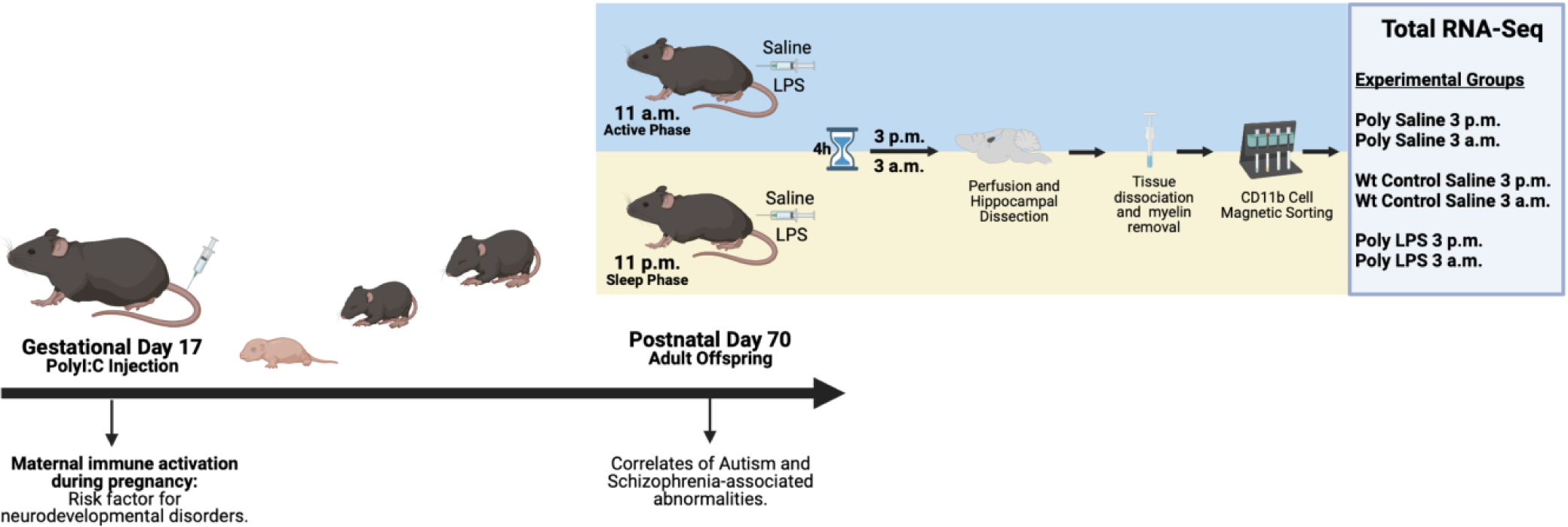
Schematic representation of the experimental setup for the transcriptional comparison of microglia from the adult offspring of the maternal immune activation (MIA) model of neurodevelopmental disorders. Maternal immune activation at gestational day 17 (GD17) was induced via injection of the synthetic viral RNA mimic Poly(I:C). The offspring was left undisturbed until they reached adulthood (postnatal day 70) and were included in the experiment. Adult offspring of PolyI:C injected dams (termed Poly animals) and controls were intraperitoneally injected with either LPS or sterile saline solution at either 11 p.m. or 11 a.m. Hippocampal microglia were isolated 4h post injection at either 3 p.m. (active phase) or 3 a.m. (sleep phase). Microglia from control animal groups that did not receive LPS (wildtype [wt] control-saline and Poly-saline), were used to investigate active and sleep phase-associated changes in transcriptional profile within Poly microglia and between Poly and control microglia at baseline. Subsequently, they were used to investigate differences in LPS response between the active and sleep phases in Poly microglia.

**Figure 7:**
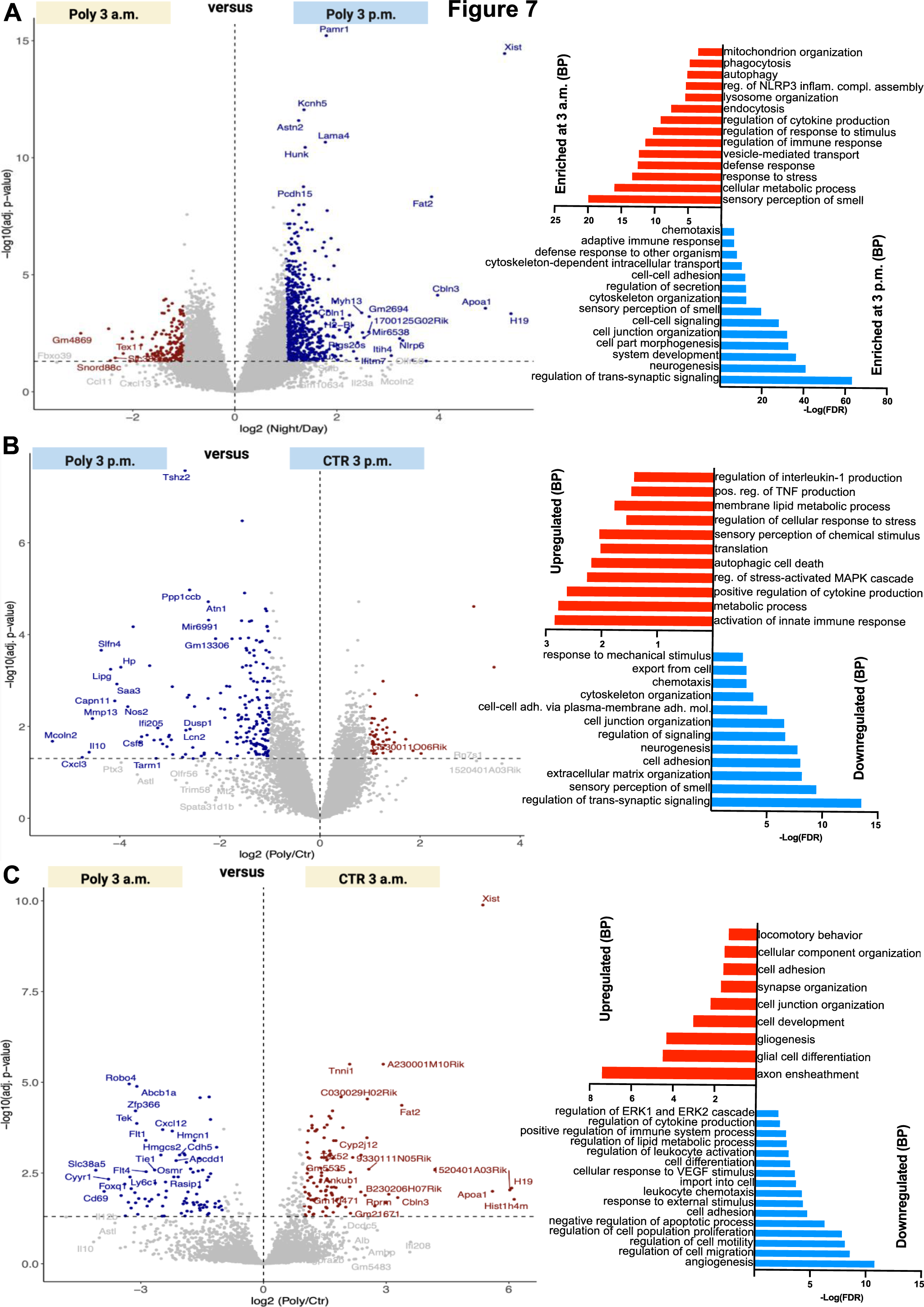
Microglia cells display a deregulated phase-associated transcriptional adaptation in the maternal immune activation model of neurodevelopmental disorders. **A** Left: Volcano plot showing the differential transcription in microglia derived from adult PolyI:C offspring (Poly animals) during the active phase (3 p.m.) versus sleep phase (3a.m.). 2527 genes were found to be differentially expressed in Poly animals between 3 p.m. and 3 a.m. (N = 3 mice/group, Log2FC cut-off of 0.5, adj. *p*-value < 0.05). **A** Right: -log(FDR) values for selected biological processes associated with the up- and downregulated genes between day and night within the Poly animal groups. **B** Left: Volcano plot representing the differential expression between microglia from Poly animals and controls during the active phase (Poly 3p.m. versus Control 3p.m.). A total of 901 genes were found to be differentially regulated (N = 3 mice/group, Log2FC cut-off of 0.5, adj. *p*-value < 0.05). **B** Right: -log(FDR) values for selected biological processes associated with the up- and downregulated genes between microglia collected during the day from Poly animals and controls. **C** Left: Volcano plot representing the differential expression between microglia isolated from Poly animals and controls during the passive phase (Poly 3a.m. versus Control 3a.m.). 250 genes were found to be differentially expressed (N = 2-3 mice/group, Log2FC cut-off of 0.5, adj. *p*-value < 0.05). **C** Right: -log(FDR) values for selected biological processes associated with the up- and downregulated genes between microglia collected during night from Poly animals and controls. Images created with Biorender.com.

### 3.6. The circadian transcriptional differences in LPS response are blunted in adult MIA offspring

Next, we sought to compare the response to LPS in Poly animals between the active and sleep phase as depicted in **Figure 6**. When comparing the LPS response in Poly microglia with the respective Poly control animals during the active phase, we found 3562 protein coding genes to be differentially expressed (1822 upregulated and 1740 downregulated, **Fig. 8A-B**, **Table S7**). LPS administration to Poly animals during the sleep phase induced a deregulation of 4601 protein coding genes (2601 upregulated and 2050 downregulated, **Fig. 8C-D**, **Table S7**). Although the upregulated biological processes were comparable, we observed an increased regulation in olfactory receptor-related biological processes amongst the downregulated genes in response to LPS during the sleep phase. To better understand the effect of time on the LPS response in Poly animals, we computed the contrast of effects (analogous to **Fig. 3D-E**) between day and night (**Fig. 8E-F**). We found that there was little interaction between the two factors, suggesting that the microglial LPS response in adult PolyI:C offspring is not affected by circadian factors as we found for control animals. Interestingly, amongst the few genes that showed an effect of time in the Poly microglial response to LPS, were the transcription factor Krüppel-like factor 4 (*Klf4*), known regulator of microglial polarization, (Kaushik et al., 2010) and *Cry1*, a core clock gene known to regulate innate immunity (**Fig. 8F**, **Table S7**).

**Figure 8:**
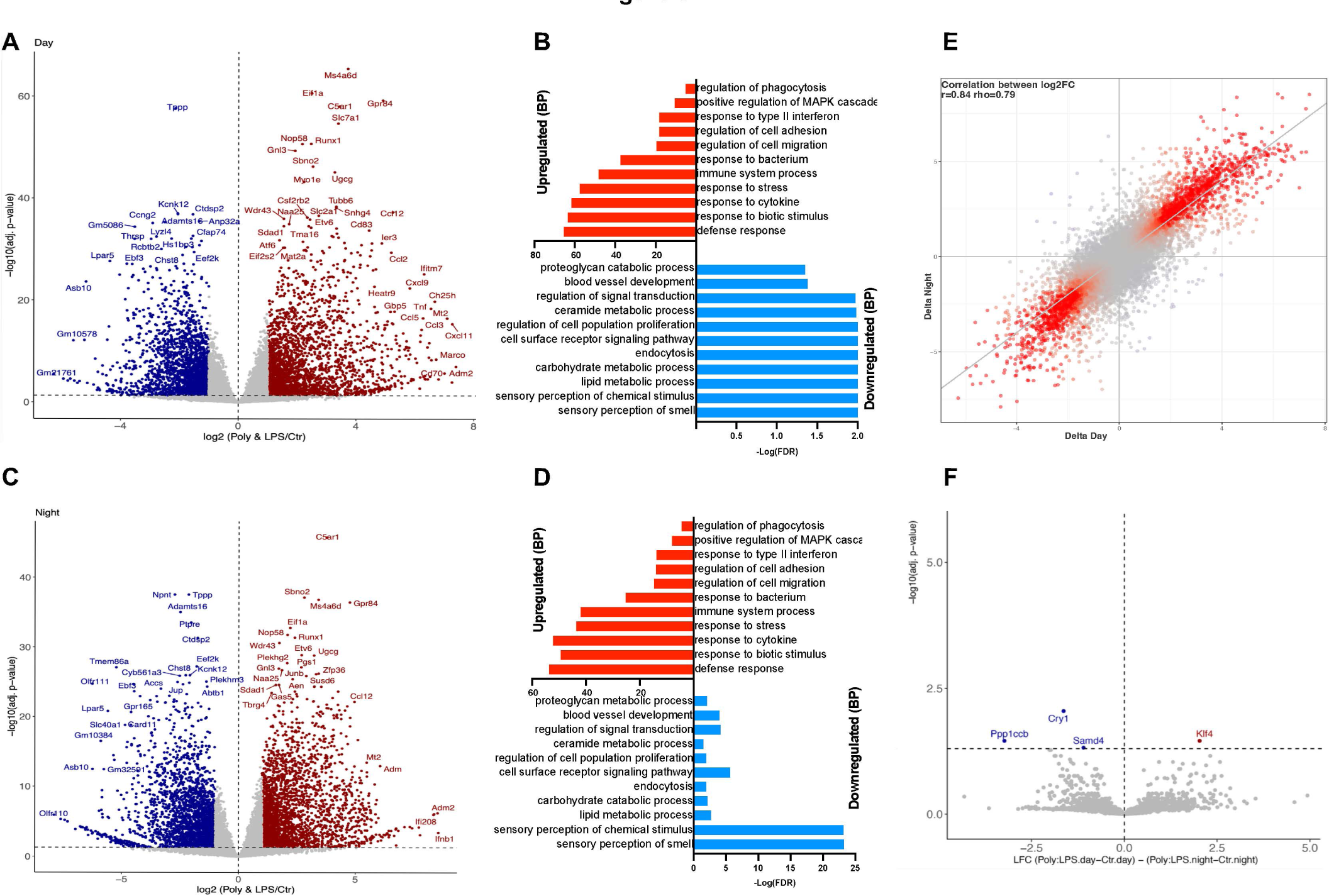
The circadian transcriptional differences in LPS response are blunted in adult MIA offspring. **A** Volcano plot showing the differential expression between microglia from Poly animals injected with LPS versus saline during the day (3 p.m., N = 3 mice/group). A total of 3562 genes were found to be differentially expressed in response to LPS administered during the day (Log2FC cut- off of 1, adj. *p*-value < 0.05). **B** -log(FDR) values for selected biological processes associated with the up- and downregulated genes in response to LPS during the day. **C** Volcano plot displaying the differential expression between microglia from animals injected with LPS versus saline during night (N = 2-3 mice/group). A total of 4601 genes were found to be differentially expressed in response to LPS administered during night (Log2FC cut-off of 1, adj. *p*-value < 0.05). **D** -log(FDR) values for selected biological processes associated with the up- and downregulated genes in response to LPS during the night. **E** Disco plot for the correlation between the absolute Log2-fold changes in gene expression in response to LPS during the active and sleep phase. **F** Volcano plot showing genes significantly regulated as an effect of time on the LPS injection (Log2FC cut-off of 1, Adj. *p*< 0.05. Images created with Biorender.com.

Based on the study design utilized in the present work, we propose an approach to account for a circadian perspective when studying microglia cells, that encompasses a given profiling within and between both control and disease groups during the active and sleep phase as illustrated in **Fig. 9**.

**Figure 9:**
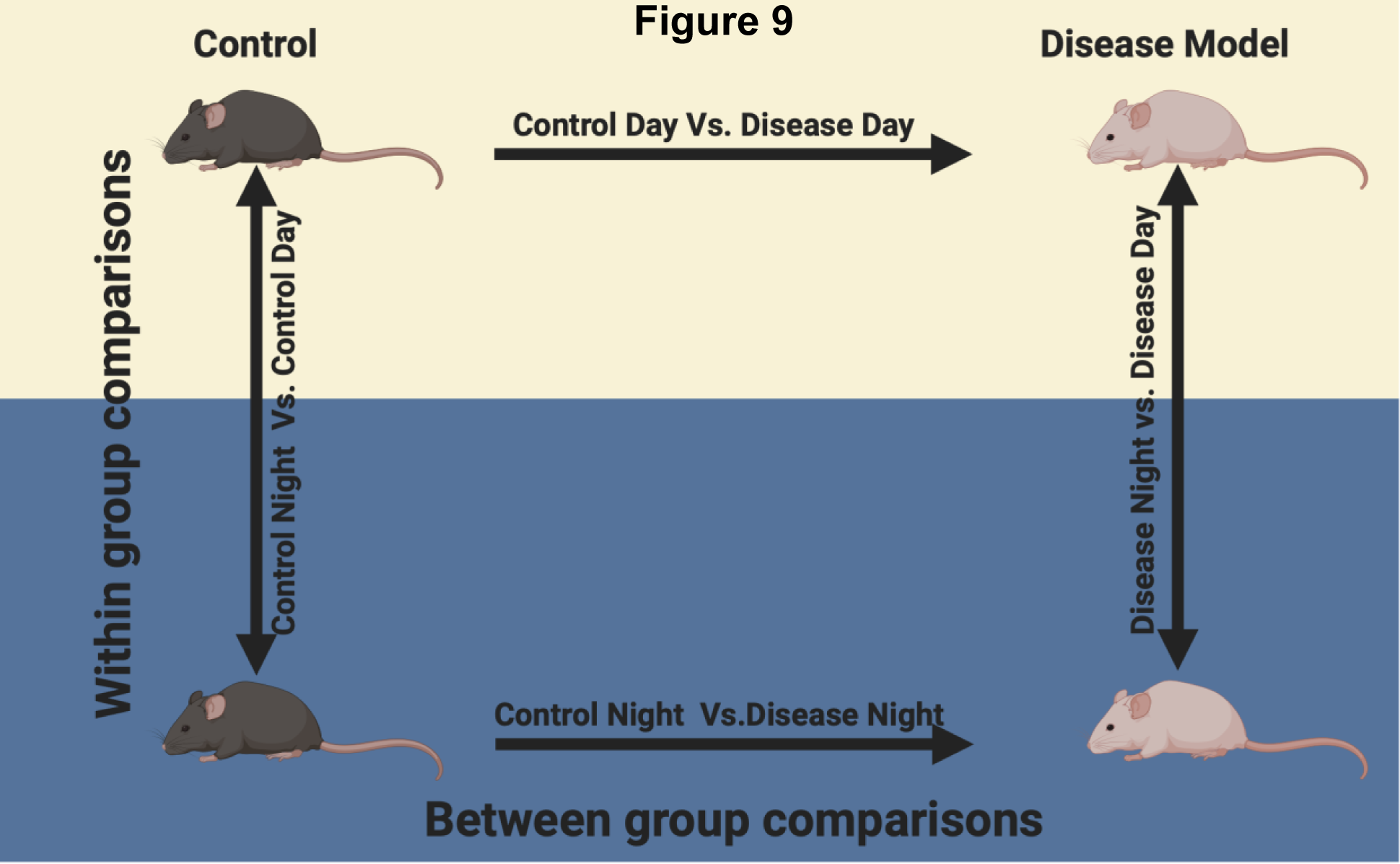
Schematic representation of the proposed general experimental design to study the microglial dynamics between the active and sleep phase in physiology (control conditions) and in pathophysiology (disease model). Image created with Biorender.com.

## 3. Discussion

Recent studies have thus far provided evidence of how manipulation of the microglial core clock genes can alter immune functions, and how microglial motility follow a circadian pattern, (Griffin et al., 2019; Stowell et al., 2019; Wang et al., 2020). However, there is at present no study directly comparing the naïve microglial transcriptional and proteotype profiles between the active and the sleep phase. Our objective was to explore differences in commonly used microglial assessments when conducted during the active phase compared to the sleep phase, thereby offering insights into whether collecting microglia at distinct circadian phases might reveal nuanced variations and provide new insights.

We observed physiological changes in genes and proteins linked to cell motility, cell adhesion, extracellular matrix remodeling, angiogenesis, and immune system processes between the light and dark phases. These alterations were mirrored by changes in functional readouts such as hippocampal synaptic pruning and intensity of response to LPS. This suggests that exclusively studying microglial functions during laboratory rodent’s sleep phase provides only a partial understanding of these cells’ physiological roles within the CNS. Our data indicate that microglia finetune their motility in a phase-dependent manner, accompanied by expression changes in neurotransmitter receptors associated with wake states, like the beta-2-adrenergic receptor (**Fig. S3A**). This has been functionally validated by two recent studies that confirmed differences in microglial process surveillance between the active and sleep phases, also in association with changes in noradrenergic tone, (Liu et al., 2019; Stowell et al., 2019). The present work provides a transcriptional and proteotype resource to understand such changes at the molecular level. Moreover, genes coding for serotonin and nicotinic acetylcholine receptors, known to modulate microglial motility and phagocytosis also showed phase-dependent regulation (**Fig. S3**), (Krabbe et al., 2012),, (Piovesana et al., 2021). This is indicative of a microglial functional adaptation to altered neurotransmitter levels between wake and sleep. Notably, we observed an active phase-associated enrichment in transcripts and proteins for the purinergic receptor P2Y12 and the enzyme CD39 (**Table S5)**, which converts extracellular ATP to ADP, the endogenous ligand for P2Y12. This system mediates the microglial process extension towards extracellular ATP sourced by e.g., high synaptic activity or brain injury (Haynes et al., 2006; Pankratov et al., 2006). Our data indicate that hippocampal microglia may be more responsive to ATP and ADP-mediated stimulation during the wake-phase, which is potentially linked to increased synaptic activity during wake and the surge in excitatory neuronal ATP occurring in the transition between sleep and wake states (Natsubori et al., 2020).

Of particular interest are the changes in expression of genes associated with ECM remodeling. We found genes coding for the ECM remodeling enzymes *Adamts4, Mmp14,* and *Ctsc*, to be altered between the sleep and active phases (**Fig. S1)**. These have been implicated in the microglial response to IL-33 for ECM-remodeling in favor of synaptic plasticity (Nguyen et al., 2020). ECM-remodeling is also of relevance in pathological conditions, e.g., in the context of brain tumors, as microglia-mediated ECM remodeling promotes brain tumor expansion, (Crapser et al., 2021). Furthermore, aberrant ECM remodeling is an important feature in neurodegenerative disorders and psychiatric conditions such as Alzheimer’s disease (AD) and schizophrenia, where abnormal ECM composition and remodeling seem to participate to glial activation and circuit instability, (Crapser et al., 2021). Furthermore, we found that genes and proteins associated with phagocytic functions were enriched during the active phase, including receptors and components of the complement system (**Fig. 2B**, **Fig. S4F**, **Table S5**). These changes were accompanied by increased hippocampal synaptic pruning during the active phase. Microglial phagocytosis is being investigated for its relevance to physiological processes like neurodevelopment, (VanRyzin, 2021) as well as in disease, such as neurodevelopmental disorders, (Lukens and Eyo, 2022; Mattei et al., 2017) and neurodegeneration, (Podleśny-Drabiniok et al., 2020). Importantly, we found that genes defining the AD-associated DAM state, (Grubman et al., 2021) are subject to phase regulation (**Table S2**). Present immune-pharmacological strategies for AD are being developed around microglial DAM-states studied mostly during the animal’s sleep (light) phase. The present study suggests that studying microglial phagocytic functions, ECM-remodeling properties as well as disease-associated states from a circadian perspective has the potential to advance our understanding of their involvement in brain disorders. This could potentially reveal circadian time-windows for optimal targeting of specific pathways. An example of such an approach is presented here for the study of microglia cells in the Poly(I:C) mediated MIA model of neurodevelopmental disorders (**Fig. 7** and **9**). We have previously shown that hippocampal microglia from adult Poly(I:C) offspring present with a transcriptional deregulation in genes associated with e.g., immune system processes, chemotaxis, and VEGF-signaling pathways, (Mattei et al., 2017). The animals used in the latter study were hosted in a light:dark cycle, and microglia were isolated during the light phase. Thus, we captured transcriptional differences between Poly(I:C) offspring and control microglia during the rodent’s sleep phase. In fact, in the present study, the comparison Poly 3 a.m. vs. Control 3 a.m. (**Fig. 7C**, sleep phase) corroborate our previous findings of downregulated pathways such as immune system processes, chemotaxis and VEGF-signaling pathways. The same comparison performed with microglia isolated during the active phase (Poly 3 p.m. vs. Control 3 p.m.) revealed an upregulation in biological processes associated with IL1 and TNF production, regulation of stress-activated MAPK cascade and metabolic processes (**Fig. 7B**).

Finally, we hypothesize that the observed microglial transcriptional, proteotype and functional changes between phases are possibly mediated by the circadian regulation in NuRD and SWI/SNF chromatin remodeling complexes (CRCs) subunits (**Table S5** and **S6**). The latter are recruited to open and close chromatin access at specific sites, and they are emerging as important players in immune cell state flexibility, (Gatchalian et al., 2020; Ramirez-Carrozzi et al., 2006). A recent study demonstrated that loss of the ARID1A subunit of the SWI/SNF complex leads to a loss of the microglial homeostatic phenotype through changes in chromatin landscape in regions that influence microglial states, (Su et al., 2022). Another study found a decrease in both NuRD and SWI/SNF subunit protein expression in aged microglia, implicating this reduction in the loss of state- polarization fluidity observed in aged microglia, (Flowers et al., 2017). We observed a protein enrichment in several CRCs subunits during the sleep phase, when microglia were associated with a stronger immune response to LPS (**Table S6**). We further noticed RNA and protein regulation in CRC subunit expression in response to LPS (**Table S6**), which may participate the microglial activation state, as shown by previous research, (Ramirez-Carrozzi et al., 2006; Roger et al., 2011). This regulation was stronger during the active phase as compared to the sleep phase, where several CRC subunits were already enriched.

### Conclusions

Investigating the circadian component in the microglial transcriptional, translational and functional programs is of high relevance for the following aspects: 1) Accounting for within and between study variability due to differences in time of sample collection and differences in animal facilities’ light schedules; 2) to gain a broader understanding of the microglial physiological roles in the brain; 3) to better understand the microglial involvement in disease and 4) to better develop pharmacological strategies to target microglia cells. Considering the present results, it would be advisable to report the lights-on/off schedules of individual animal facilities, and the specific hour post light-on (or off) when animals are euthanized for microglial characterizations. This may support a better reproducibility of scientific findings. For the purpose of implementing a circadian component in microglial characterizations, cohorts of mice can be hosted in rooms set in a dark:light and in a light:dark cycle in parallel. This way, cohorts would be available during both phases, facilitating experimental designs such as the one proposed in **Fig. 9**. A limitation of the present study is that we did not perform a complete circadian transcriptional time-course, but rather had fixed time-points in the light and dark phase. Collins et al. recently demonstrated how the transcriptome and proteome of peripheral macrophages undergoes circadian oscillations, which influences their immune responsiveness and phagocytic functions, similar to what we found in microglia cells in the present study, (Collins et al., 2021b) Moreover, we only performed this analysis in male mice. Future studies should aim at addressing how the microglial transcriptional and translational profiles changes throughout a complete circadian cycle and in a sex- dependent manner.

## 4. Material and methods

### 4.1. Animals

Nine to ten weeks old male C57BL6/N mice (Charles River Laboratories, Sulzfeld, Germany) were used throughout the study. They were caged 3-5 animals per cage in individually ventilated cages (IVCs). The animal vivarium was a specific-pathogen-free (SPF) holding room, which was temperature- and humidity-controlled (21 ± 3 °C, 50 ± 10%) and kept under a reversed light–dark cycle (lights off: 09:00 AM–09.00 PM). Hence, all animals were in their active and sleep phases during the experimenters’ light (day) and dark (night) phases, respectively. All animals had *ad libitum* access to food (Kliba 3436, Kaiseraugst, Switzerland) and water throughout the entire study. All procedures described in the present study had been previously approved by the Cantonal Veterinarian’s Office of Zurich, and all efforts were made to minimize the number of animals used and their suffering.

### 4.2. Breeding and maternal immune activation

The maternal immune activation (MIA) model was implemented as previously described (Scarborough et al., 2021). Briefly, successful mating was verified by the presence of a vaginal plug, considered as gestational day 0 (GD 0). Dams were housed individually throughout gestation. On GD 17, pregnant dams were randomly assigned to a single intravenous tail-vein injection of either poly(I:C) (Sigma–Aldrich P9582, Buchs, St. Gallen, Switzerland) to induce MIA, or pyrogen-free 0.9% NaCl (prenatal control). We previously ascertained the quality, molecular composition and immunopotency of the poly(I:C) batch (#086M4045V) used in this study (Mueller et al., 2019). Based on our previous dose–response studies (Mueller et al., 2018)and molecular composition of the poly(I:C) batch, poly(I:C) was administered intravenously (i.v.) into the tail vein at a dose of 5 mg/kg. The gestational time point of poly(I:C) or vehicle administration (i.e., GD 17) was selected based on previous studies showing more extensive abnormalities in hippocampal structures and functions when MIA occurs in late gestation as compared to earlier time points (Giovanoli et al., 2015; Meyer et al., 2008, 2006). All injections had a total volume of 5 ml/kg. Following the injection, the dams were placed back in their home cages and left undisturbed until the first cage change on postnatal day 7. Offspring were weaned on postnatal day 21 and littermates of the same sex were caged separately and maintained in groups of 4–5 animals per cage.

### 4.3. *In vivo* LPS treatment

LPS (Enzo Lifesciences, serotype O55:B5, cat. Nr. ALX-581-013-L002) was diluted in sterile saline solution and injected intraperitoneally at a concentration of 1mg/kg. Control animals were injected with saline solution. To avoid batch to batch effects on the microglial immune response, all experiments with LPS were performed within 24h using the same batch of LPS.

### 4.4. Brain dissociation and cell isolation

Brain tissue dissociation and microglia cell isolation were performed according to a recently optimized mechanical dissociation (MD) protocol (Mattei et al., 2020). The protocol is carried out at 4°C to minimize microglial cell activation due to the isolation procedure (Mattei et al., 2020). Briefly, the animals were deeply anesthetized with an overdose of Nembutal (Abbott Laboratories, North Chicago, IL, USA) and transcardially perfused with ice-cold, calcium- and magnesium-free Dulbecco’s phosphate-buffered saline (DPBS). Hippocampi and frontal cortices were dissected on a cooled petri dish and placed in ice-cold Hibernate-A medium. MD at 4°C was carried out on ice with a Dounce homogenizer, using a loose pestle. Myelin debris were removed via percoll gradient centrifugation. After washing of the percoll, total brain cell pellets were used for CD11b magnetic-activated cell sorting (MACS) using anti-mouse CD11b magnetic microbeads (Miltenyi) according to the manufacturer’s instructions. The MACS buffer consisted of 3% bovine serum albumin (BSA) diluted in DPBS from a 7.5% cell-culture grade BSA stock (Thermo Fisher Scientfic). Total hippocampal cell pellets were re- suspended in 90 μl MACS buffer and 10 μl anti-mouse-CD11b magnetic beads. The cells were then incubated for 15 min at 4°C. Cells were washed with 1 ml MACS buffer. The cells were then passed through an MS MACS column (Miltenyi) attached to a magnet. After washing the columns three times with MACS buffer, microglia were flushed from the column with 1 ml MACS buffer and pelleted. Cell pellets were then snap-frozen in liquid nitrogen and stored at -80°C for further analysis.

### 4.5. Flow cytometry-based intracellular detection of synaptic particles in microglia

We used a recently established protocol (Brioschi et al., 2020) to detect and quantify the synaptic marker, vesicular glutamate transporter 1 (vGLUT1) in microglia by flow cytometry. Following percoll isolation (as described above, section 2.4), total cell pellets were sequentially stained with fixable live/dead staining (1/1000 in PBS, ThermoFisher, #L34969, 30 min) and for the microglial surface markers CD11b (1/100, ThermoFisher Scientific, #25-0112-82), CD45 (1/100, BD Biosciences, #559864), Ly6C (1/100, BD Biosciences, #553104) and Ly6G (1/100, BD Biosciences, #551460), and CD16/CD32(1/200, ThermoFisher Scientific, #14-0161-82). Subsequently, stained samples were fixed and permeabilized using the BD Cytofix/Cytoperm kit according to manufacturer’s instructions (#554714). Following fixation, intracellular staining for vGLUT1 (1/200, Miltenyi Biotec, #130-120-764, 1h in 1x BD Permeabilization Buffer) was immediately performed. Samples were acquired on BD FACS Fortessa (BD Bioscience) flow cytometer and microglia population was defined as CD11b^++^/ CD45^+^/ Ly6C^-^/ Ly6G^-^/ viable cells. Splenocytes were used as negative control to set the threshold and gate on vGLUT1-positive microglia fractions. Raw data were analyzed with FlowJo v10 (BD Bioscience).

### 4.6. RNA extraction and quantification

The RNA was extracted via the Lexogen Split-RNA extraction kit (cat. number 008.48) according to the manufacturer’s instructions. RNA concentrations were measured via Qubit 4 fluorometer (Invitrogen), using the RNA HS Assay kit (Invitrogen).

### 4.7. Total RNA library preparation and sequencing

Before library preparation, RNA integrity was assessed on an Agilent TapeStation system 4150 using the RNA screen tape (Agilent). 50 ng total RNA were used as input for ribosomal RNA (rRNA) depletion using the NEBNext rRNA depletion kit (New England BioLabs inc., product code: E6350) according to the manufacturer’s instructions. Following rRNA depletion, total RNA libraries were built using the NEBNext Ultra II library prep kit for Illumina (New England BioLabs inc., product code: E7775) according to the manufacturer’s instructions. The yield of amplified libraries was measured on a Qubit 4 fluorometer using the Qubit high sensitivity DNA kit (HS DNA kit). Amplified libraries were further analyzed on the HS D1000 screen tape on a TapeStation system 4150 to assess library size and molarity prior to pooling. The libraries were sequenced using Illumina HISeq-4000 at a depth of 40 million reads/sample.

### 4.8. Computational analysis of RNA-sequencing data

RNA-Seq reads were mapped to the mouse genome (GRCm38, version M12 (Ensembl 87)) with STAR (Dobin et al., Bioninfomatics 2013, version-2.7.3a) using the following parameters –outSAMunmapped Within --outFilterType BySJout -- outFilterMultimapNmax 20 --alignSJoverhangMin 8 --alignSJDBoverhangMin 1 --outFilterMismatchNmax 999 --outFilterMismatchNoverLmax 0.04 --alignIntronMin 20 -- alignIntronMax 1000000 --alignMatesGapMax 1000000. Reads were assigned to genes using FeatureCounts (Liao et al., Bioinformatics 2014, version-v2.0.0) with the following parameters: -T 2 -t exon -g gene_id -s 0. For the differential expression analyses we used DESeq2 (version-1.32.0) with default parameters (Love et al., 2014). We filtered genes that had less than 5 counts in at least 3 samples. The model.matrix design was specified as “∼0 + group”, where “group” variable was defined as a combination of time and treatment (ie. ctr.day [active phase], ctr.night [sleep phase], poly(I:C).day [active phase], poly(I:C).night [slpee phase], LPS.day [active phase], LPS.night [sleep phase], LPS. poly(I:C).day [active phase], LPS. poly(I:C).night [sleep phase]). The gene set enrichment analyses were carried out using CERNO algorithm from R tmod package (version-0.50.06), (Zyla et al., 2019) and the Panther (Mi et al., 2013) online tool.

### 4.9. Quantitative real-time polymerase chain reaction (qRT-PCR)

RNA was analyzed by TaqMan qRT-PCR instrument (CFX384 real-time system, Bio-Rad Laboratories) using the iTaq™ Universal Probes One-Step Kit for probes (Bio-Rad Laboratories). The samples were run in 384-well formats in triplicates as multiplexed reactions with a normalizing internal control. We chose *36B4* as internal standard for gene expression analyses. Thermal cycling was initiated with an incubation at 50°C for 10 min (RNA retrotranscription) and then at 95°C for 5 min (TaqMan polymerase activation). After this initial step, 39 cycles of PCR were performed. Each PCR cycle consisted of heating the samples at 95°C for 10 s to enable the melting process and then for 30 s at 60°C for the annealing and extension reaction. Cutsom-made primers with probes for TaqMan were purchased from Thermo Fisher: housekeeping gene: *36B4*, product code: NM_007475.5; *sialic acid-binding immunoglobulin-like lectin-H (Siglech)*, product code: Mm_00618627_m1: *purinergic receptor P2Y12* (*P2ry12)*, product code: Mm00446026_m1; *purinergic receptor P2Y6 (P2ry6)*, product code Mm02620937_s1; and *transcription factor PU.1 (Spi1)*, product code Mm00488140_m1. Relative target gene expression was calculated according to the Delta C(T) method.

### 4.10. Quantification of cytokines and fractalkine

Animals were deeply anesthetized with an overdose of Nembutal (Abbott Laboratories, North Chicago, IL, USA) and transcardially perfused with 15 ml ice-cold DPBS.

Hippocampi were quickly removed, snap-frozen in liquid nitrogen and stored at -80°C. Tissue lysis was performed in Roche cOmplete Lysis-M reagents (lysis buffer and protease inhibitor cocktail contained in the kit, Millipore Sigma cat. Nr. 4719956001). Hippocampi were placed in protein low-binding tubes mounted onto a Qiagen TissueLyser II (Qiagen cat. Nr. 85300). Lysates were span down at 20.000 rcf for 20 minutes at 4°C to remove debris, and the supernatant was stored at -80°C.

Cytokine protein levels in hippocampal lysates were quantified using a Meso-Scale Discovery (MSD) V-Plex electrochemoluminescence assay for mice according to the manufacturer’s instructions (V-Plex proinflammatory panel 1 mouse kit, MSD cat. Nr. K15048D-1). The assay allowed for the detection and quantification of interleukin (IL)-1β, IL-6, IL-4, IL-10, and tumor necrosis factor (TNF)-α . Samples were run in duplicate, and the plate was read via a SECTOR PR 400 (MSD) imager and analyzed using MSD’s Discovery Workbench analyzer and software package. Fractalkine protein levels were measured in hippocampal lysates via the R&D System’s Mouse Fractalkine (CX3CL1) Quantikine ELISA Kit (R&D Systems cat. Nr. MCX310) according to the manufacturer’s instructions. Samples were measured in duplicate, and the plate was read on a microplate reader (SPARK® multimode microplate reader, Tecan Trading AG, Switzerland). Cytokines and fractalkine concentrations were normalized to the total protein amount in each sample.

### 4.11. Proteotype analysis

#### 2.11.1. Liquid chromatography–tandem mass spectrometry (LC–MS/MS) analysis

For MS analysis, peptides were reconstituted in 5% acetonitrile and 0.2% formic acid containing standard iRT peptides (Biognosys AG, Switzerland). The peptides were analyzed in data-independent acquisition (DIA) mode and data-dependent acquisition (DDA) mode for spectral library generation. For spectral library generation, a fraction of the samples originating from the same condition were pooled to generate a mixed sample for each condition. Peptides were separated by reverse-phase chromatography on an 25-cm Easy Spray column (Thermo Fisher Scientific, USA) connected to an EASY-nLC 1200 instrument (Thermo Fisher Scientific). The HPLC was coupled to a Q Exactive HF mass spectrometer equipped with a nanoelectrospray ion source (Thermo Fisher Scientific). For each injection, approximately 2 µg peptides were loaded onto the column and separated via a 180 min gradient from 100% buffer A (99% H2O, 0.1% formic acid) to an increasing percentage of buffer B (99.9% acetonitrile, 0.1% formic acid). The DIA method contained 24 DIA segments with varied m/z widths over a mass range of 350 to 1650 m/z with 30,000 resolution. Injection time (IT) was set to auto, an automatic gain control (AGC) of 3 x 106 was applied, and a survey scan of 120,000 resolution (54 ms maximum IT, 3 x 106 AGC target; default charge state of 3, loop count of 1, and 27 NCE [1]). For the DDA, a Top10 method was recorded with 60,000 resolution of the MS1 scan (54 ms max IT, AGC target of 3 x 106), followed by high-energy collisional dissociation (HCD)-MS/MS scans at 15,000 resolution of the MS1 scan (54 ms max IT, AGC of 5 x 104). To avoid multiple scans of dominant ions, the precursor ion masses of scanned ions were dynamically excluded from MS/MS analysis for 30 s. Single-charged ions and ions with unassigned charge states or charge states above 6 were excluded from MS/MS fragmentation. The covered mass range was identical to the DIA.

#### 2.11.2. Data analysis DIA LC-MS/MS

LC-MS/MS DIA runs were analyzed using a spectral library generated from the pooled samples measured in DDA. The collected DDA spectra were searched against UniprotKB (UniProt Swissprot, Mus musculus retrieved in 2018) using the Sequest HT search engine within Thermo Proteome Discoverer (PD) version 2.3 (Thermo Fisher Scientific). The following PD parameters were applied: two missed cleavages, specific tryptic digestion, fixed carbamidomethylation of C and oxidation of M, variable deamidation of

R and acetylation at the N-terminus. Monoisotopic peptide tolerance was set to 10 ppm, and fragment mass tolerance was set to 0.02 Da. The identified proteins were assessed using Percolator and filtered using the high peptide confidence setting in PD. Identification results were imported to Spectronaut Pulsar version 14.0 (Biognosys AG) for the generation of spectral libraries. Targeted data extraction of DIA-MS acquisitions was performed with Spectronaut with default settings using the generated spectral libraries.

Extracted features were exported from Spectronaut for statistical analysis with MSstats (version 3.18.0) using default settings, and iRT peptides to normalize across runs. Features were filtered for calculation of Protein Group Quantity as defined in Spectronaut settings; common contaminants were excluded. In MSstats, the model estimated fold change (FC) and statistical significance for all compared conditions. Significantly different proteins were determined by an adjusted *p*-value < 0.05 for the comparisons of control conditions (control 3 a.m. versus control 3 p.m.). For the differential expression in response to LPS the threshold was set at Log2-fold-change > 1 and adjusted *p*< 0.05. Benjamini-Hochberg method was used to account for multiple testing.

### 2.12. Statistical analysis

All statistical analyses of ELISA, electrochemiluminescence, qRT-PCR and flow cytometry data were performed using Graphpad Prism (Version 10.0.2; GraphPad Software, La Jolla, California), with statistical significance set at *p* < 0.05. Group-wise comparisons were analyzed using independent Student’s *t* tests (two-tailed). For the transcriptional analysis of microglia between 1 a.m. and 1 p.m., and the transcriptional analysis of microglia from Poly(I:C) offspring at baseline, differentially expressed genes (DEGs) were defined by a Log2-fold change (Log2FC) cut-off of 0.5 and an adjusted (adj.) *p*-value cut-off of 0.05. DEGs presented in heatmaps in Fig.S1-S4 include genes that were up- or down-regulated based on an adj. *p*-value of 0.05. For the transcriptional and proteotype analysis of the LPS responses, DEGs were defined by a Log2FC cut-off of 1 and adj. *p*-value of 0.05. For the transcriptional and proteotype comparison of microglia cells between 3 a.m. and 3 p.m., since the transcriptional changes could be validated at the proteotype level even for minor Log2FCs, differentially expressed genes and proteins were defined by an adj. *p*-value cut-off of 0.05.

## Author Contributions

D.M., A.I., D.B., S.A.W., B.W., H.K. and U.M. conceived and designed the study. D.M., A.I., J.H., P.P., B.U., S.S. J.R., U. W.-S., and J.S. performed the experiments and analyzed the data. D.M., A.I. and U.M. wrote the manuscript. All authors discussed the results and commented on the manuscript.

## Data Availability

All mass spectrometric data and acquisition information will be deposited to the ProteomeXchange Consortium (www.proteomexchange.org/) via the PRIDE partner repository (Perez-Riverol et al., 2019) at the time of publication.

The raw RNA-sequencing data will be deposited in NCBI’s Gene Expression Omnibus and will be accessible through GEO Series accession number at the time of publication.

## Acknowledgments

The authors would like to thank Tarek Khashan and Dr. Ester Del Duca for technical support in generating the figures.

---

Table S1: Complete table for the differentially expressed microglial genes (DEGs) between 1 p.m. and 1 a.m. followed by gene ontology terms for biological processes associated with the DEGs. All genes differentially expressed (adj. *p*< 0.05) are included in the table.

Table S2: Table displaying the differential gene expression of key homeostatic and disease-associated microglial (DAMs) genes between 1 a.m. and 1 p.m.

Table S3: Complete table for the differentially expressed microglial genes and proteins in response to LPS at 3 p.m. (Day) and 3 a.m. (Night) followed by gene ontology terms for biological processes associated with the deregulated genes and proteins in each condition. All genes differentially expressed (adj. *p*< 0.05) are included in the table.

Table S4: Complete table for the differentially expressed microglial genes and proteins between 3 p.m. (Day) and 3 a.m. (Night) followed by gene ontology terms for biological processes associated with the DE genes and proteins. All proteins differentially expressed (adj. *p*< 0.05) are included in the table.

Table S5: Table showing transcriptional and proteotype changes in genes and proteins relevant for microglial mitochondrial and metabolic functions, immune activation, motility and phagocytosis, extracellular matrix components and chromatin remodeling. Displayed are the Log2 fold changes (Log2FC) and adjusted *p*-values for the differential gene expression between 1 a.m. and 1 p.m., 3 a.m. and 3 p.m. followed by the changes in protein expression between 3 a.m. and 3 p.m. Positive Log2FCs indicate an enrichment in the sleep phase (1 a.m. and 3 a.m.), negative Log2FCs indicate an enrichment in the active phase (1 p.m. and 3 p.m.).

Table S6: Table showing transcriptional and proteotype changes between 3 a.m. and 3 p.m. in genes and proteins responsible for the epigenetic remodeling of chromatin accessibility at baseline (Green) and in response to LPS at either 3 p.m. (Day, orange) or 3 a.m. (Night, blue). Displayed are the Log2 fold changes (Log2FC) and for the differential gene/protein expression between 3 a.m. and 3 p.m. Positive Log2FCs indicate an enrichment in the sleep phase (Night), negative Log2FCs indicate an enrichment in the active (Day) phase. Epigenetic writers (bottom) are enzymes responsible for the acetylation and methylation of histones and DNA. Epigenetic readers are proteins able to recognize and bind to specific histone and DNA epigenetic marks. Epigenetic readers such as BRDs can recruit SWI/SNF complex subunits, while CHDs can be integral parts of the NuRD complex. Further displayed are key subunits of the main chromatin remodeling complexes. Several components of the chromatin remodeling machinery are enriched in the sleep phase (3 a.m., green). In response to LPS there is a marked regulation in RNA and protein expression of the chromatin remodeling machinery subunits. This regulation appears to be stronger during the active phase (3 p.m., Day), likely because several components are already enriched during the sleep phase.

Table S7: Complete table for the differentially expressed microglial genes in adult offspring of the PolyI:C maternal immune activation model of neurodevelopmental disorders, at baseline and in response to LPS at 3 p.m. (Day) and 3 a.m. (Night) followed by gene ontology terms for biological processes associated with the deregulated genes in each condition. All genes differentially expressed (adj. *p*< 0.05) are included in the table.

## Supplementary Figures

**Supplementary Figure S1:**
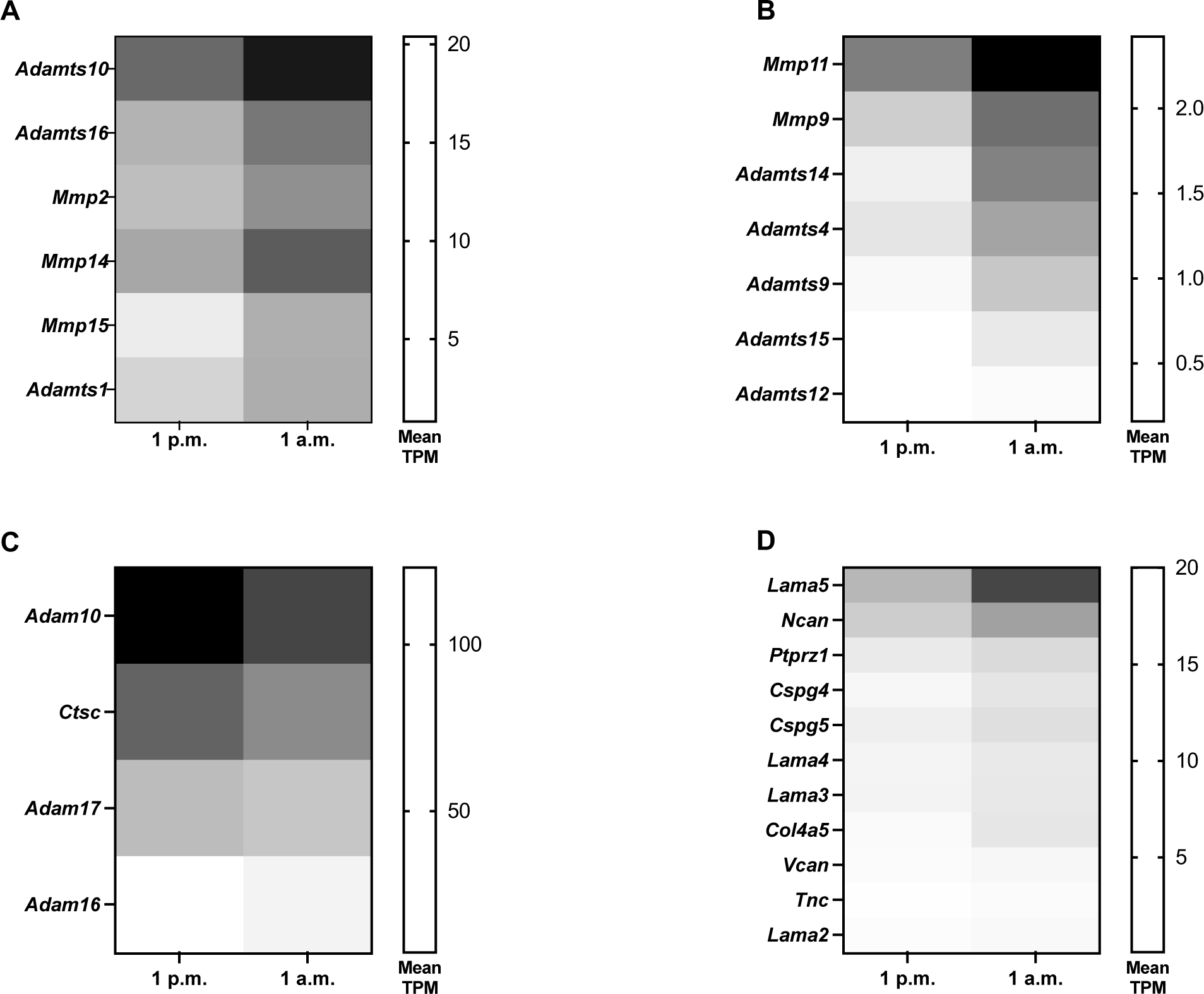
Microglial genes associated with extracellular matrix remodeling are altered between the active and sleep phases. A-D Heatmaps displaying the changes in average transcripts per million (TPM) for genes coding for enzyme dedicated to the extracellular matrix (ECM) breakdown A-C and genes coding for ECM components D in the microglial transcriptional comparison between 1 a.m. and 1 p.m. Only genes that were significantly differentially expressed are displayed (adj. *p*< 0.05).

**Supplementary Figure S2:**
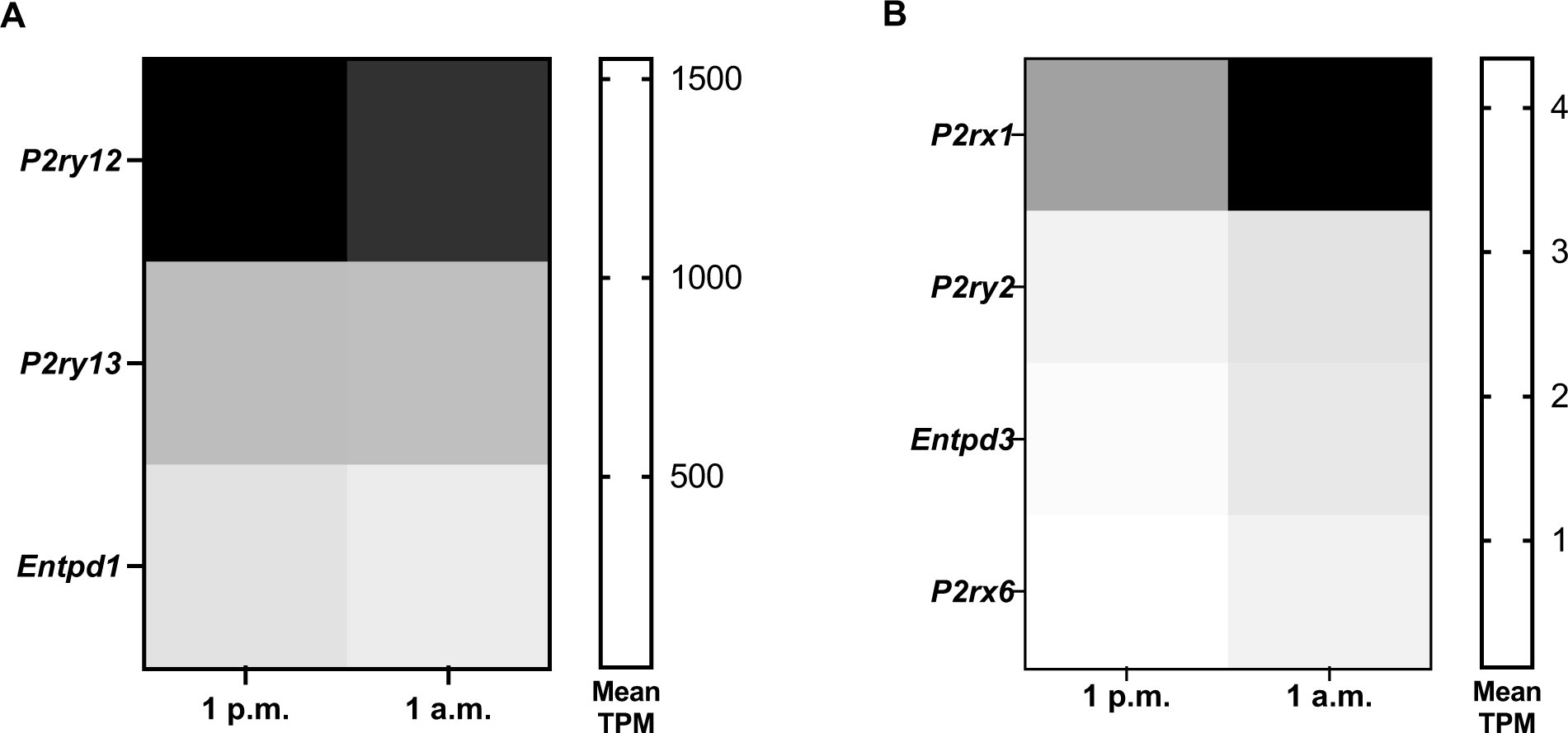
Genes associated with the microglial purinergic signaling are altered between the active and aleep phase. Heatmaps displaying the changes in average transcripts per million (TPM) for highly A and lowly B expressed genes coding for purinergic receptors and enzymes involved in the breakdown of extracellular nucleotides in the microglial transcriptional comparison between 1 a.m. and 1 p.m. Only genes that were significantly differentially expressed are displayed (adj. *p*< 0.05).

**Supplementary Figure S3:**
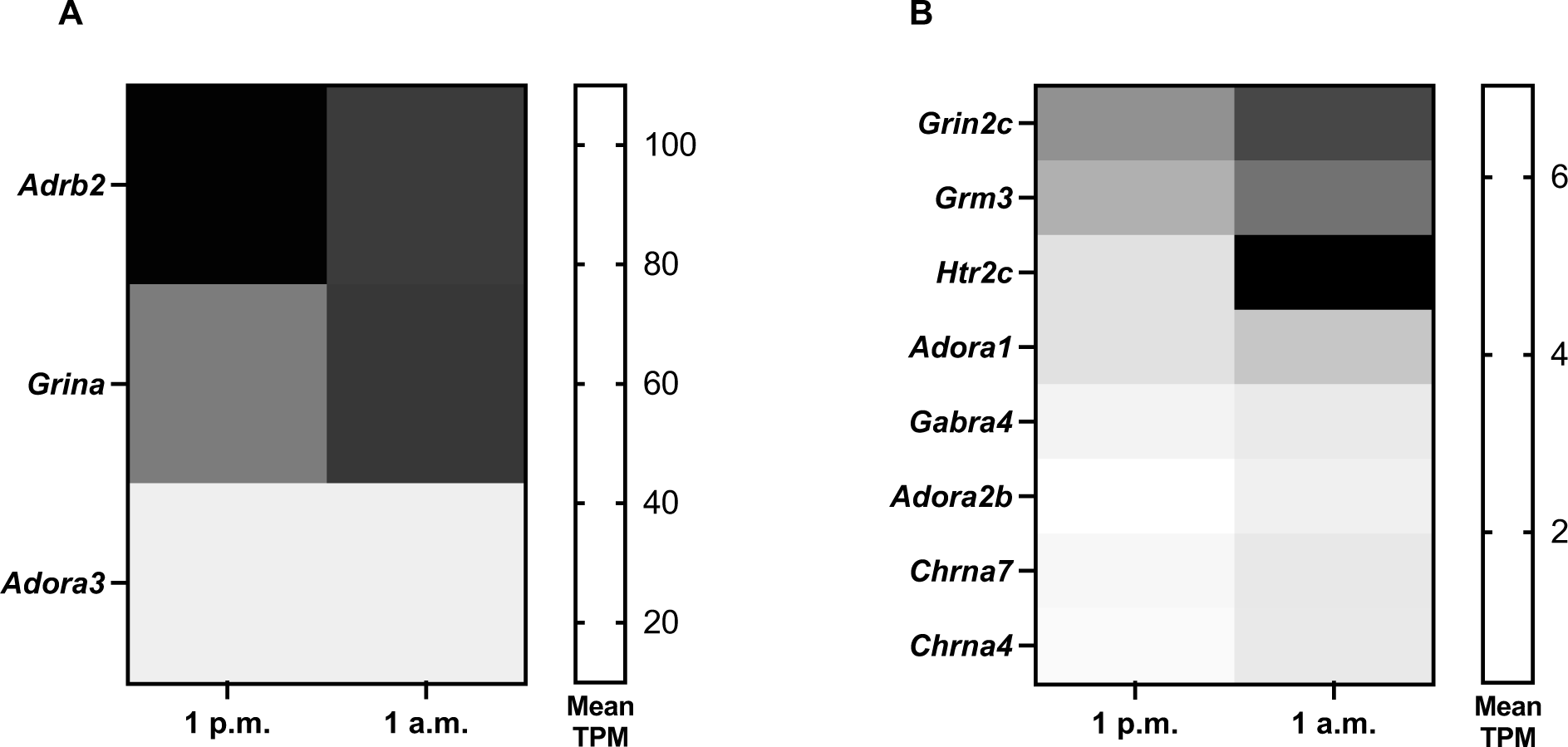
The microglial expression of neurotransmitter receptors is altered between the active and sleep phase. Heatmaps displaying the light- and dark-phase associated changes from the microglial transcriptional comparison between 1 a.m. (Night) and 1 p.m. (Day), in average transcripts per million (TPM) for highly A and lowly B expressed genes coding for neurotransmitter receptors. Only genes that were significantly deregulated are displayed (adj. *p*< 0.05).

**Supplementary Figure S4:**
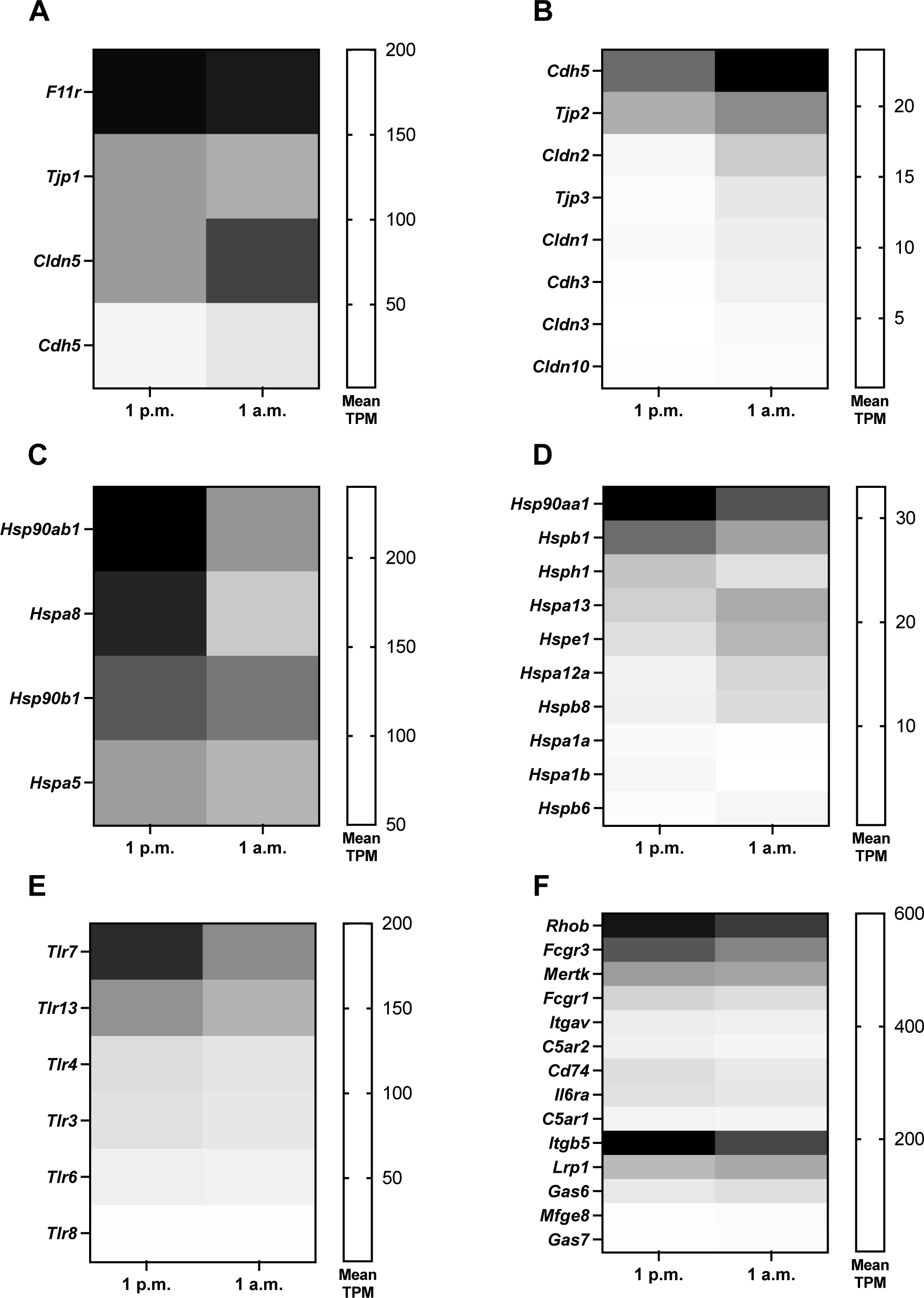
Active and sleep phase-associated transcriptional changes in genes associated with microglial key functions. A-D Heatmaps displaying the active and passive phase-associated changes from the microglial transcriptional comparison between 1 a.m. and 1 p.m., in average transcripts per million (TPM) for highly A and lowly B expressed genes associated with cell-to-cell adhesion and junction formation and heat-shock proteins C-D. E Heatmap showing the active and passive phases-associated differences in mean TPM for genes coding for Toll-like receptors. F Heatmap displaying the differential expression in genes relevant for microglial activation and phagocytosis. Only genes that were significantly deregulated are displayed (adj. *p*< 0.05).

**Supplementary Figure S5:**
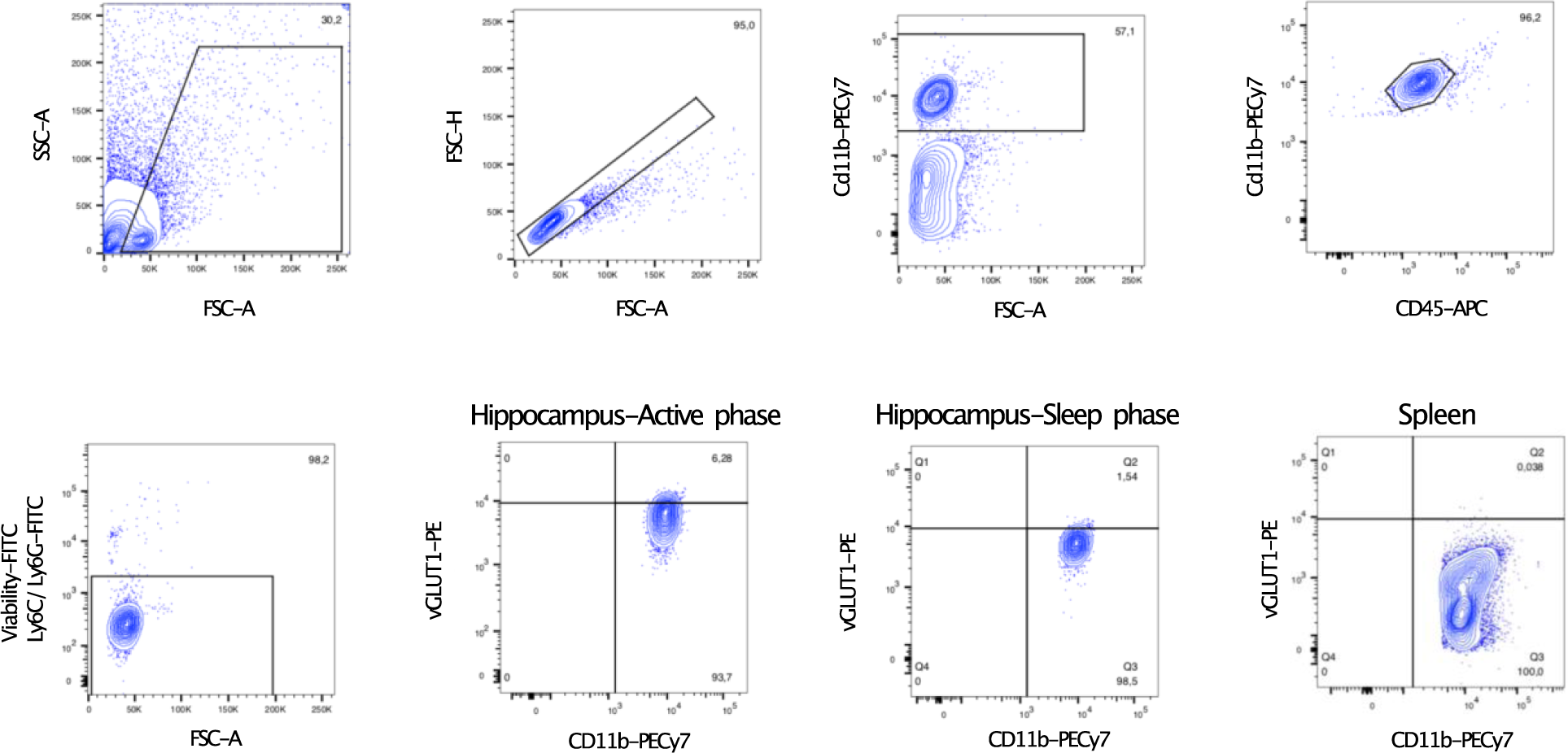
Gating strategy for the flow cytometry analysis of vGLUT1- positive inclusions within hippocampal microglia cells sorted during wither the active or the sleep phase. Supplementary Figure S6: A -log(FDR) values for biological processes associated with the genes deregulated in response to LPS in function of time of injection. B Unbiased Tmod enrichment analysis for reactome gene sets for the LPS response genes regulated by time of injection.

**Supplementary Figure S6:**
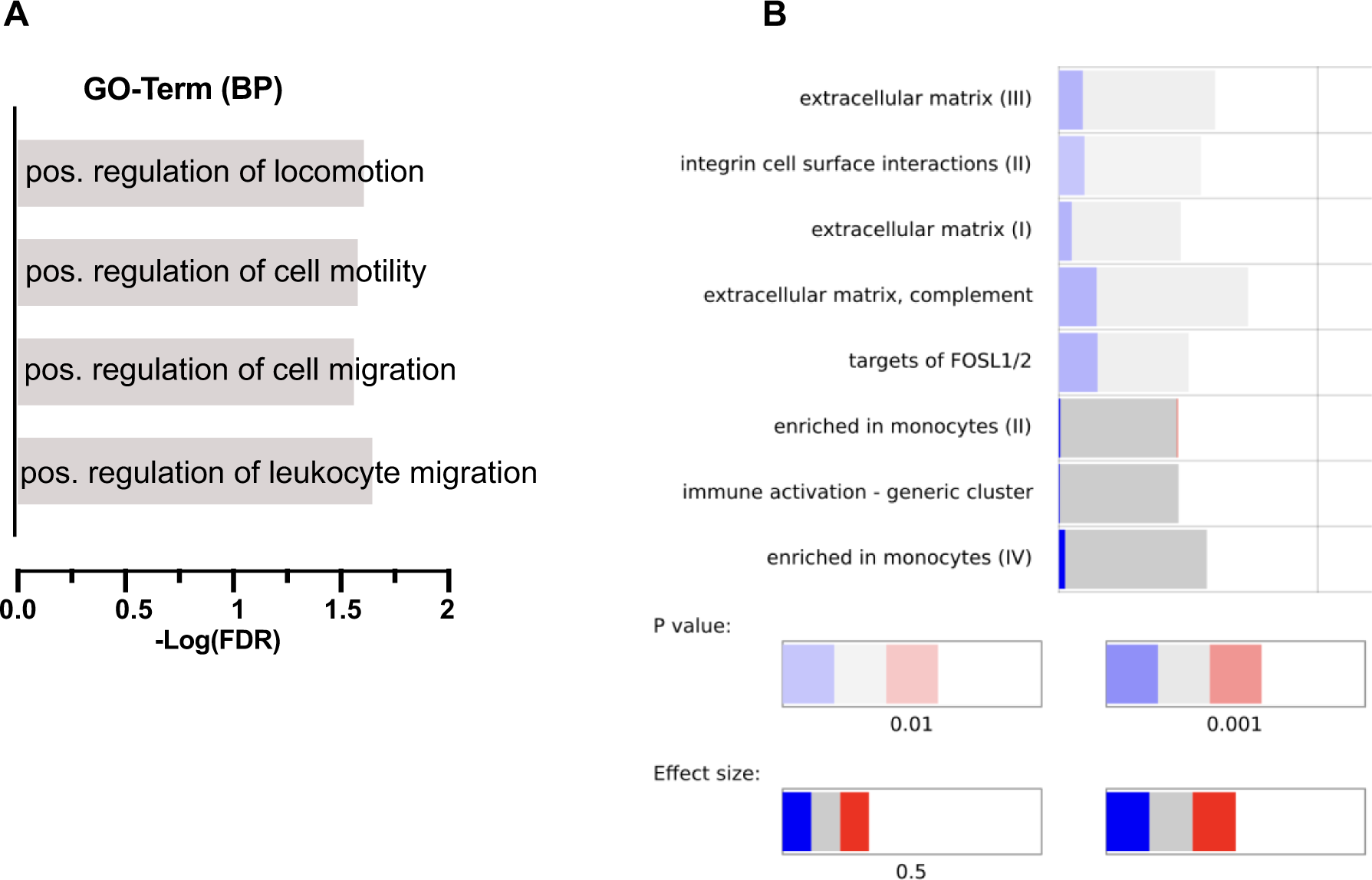
A -log(FDR) values for biological processes associated with the genes deregulated in response to LPS in function of time of injection. B Unbiased Tmod enrichment analysis for reactome gene sets for the LPS response genes regulated by time of injection.

**Supplementary Figure S7:**
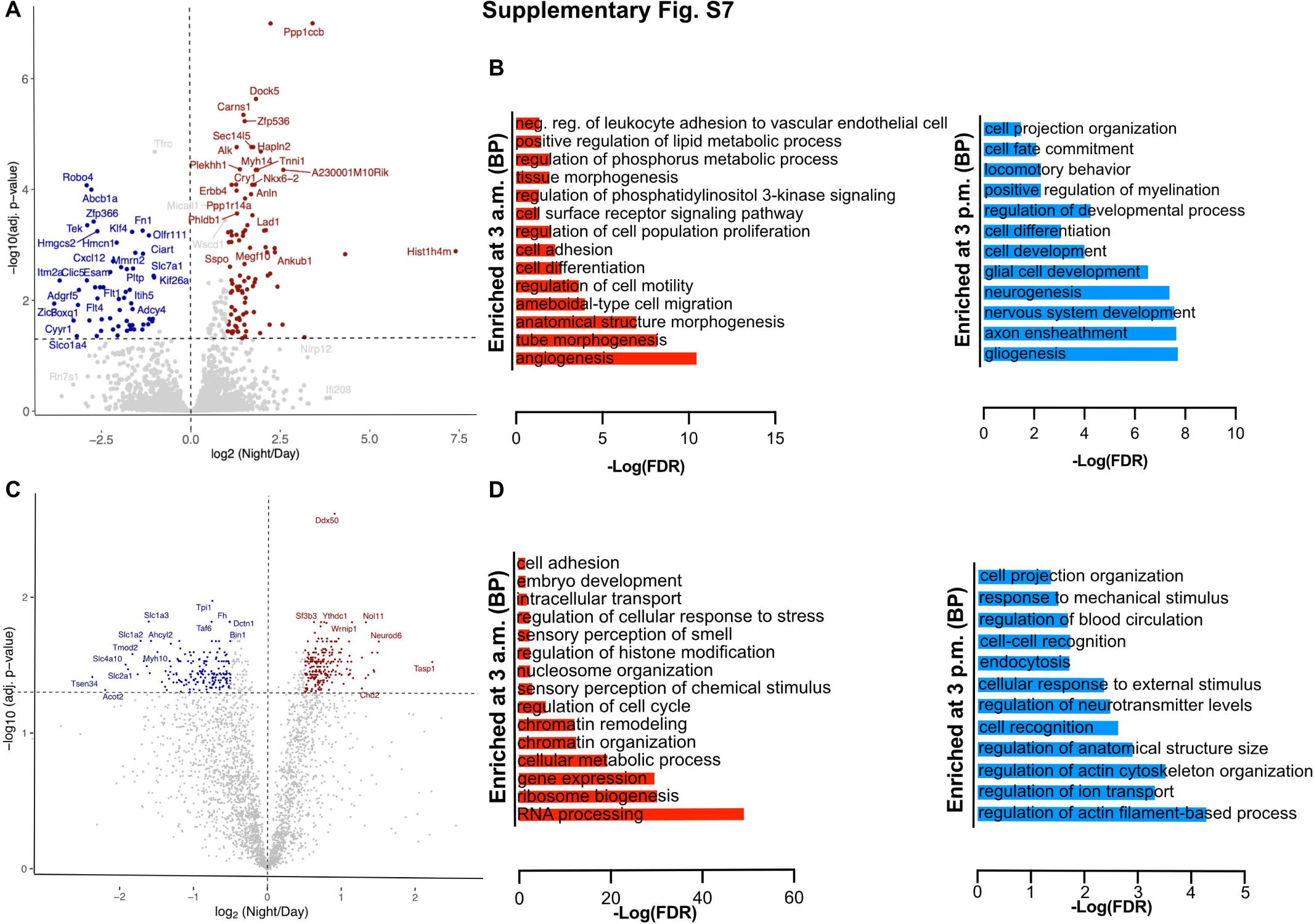
The microglial transcriptional and proteotype regulation between the active (3p.m.) and sleep (3 a.m.) phases. A Volcano plot showing the differential gene expression in microglia between 3 a.m. (sleep phase) and 3 p.m. (active phase). 182 genes were found to be differentially expressed (adj. *p*< 0.05, N = 2-3/group). B -Log(FDR) values for selected biological processes from the gene ontology analysis of the up- and downregulated genes between 3 a.m. and 3 p.m. C Volcano plot displaying the differentially expressed proteins between 3 a.m. (sleep phase) and 3 p.m. (active phase) 481 proteins were found to be differentially regulated between the light and dark phase (adj. *p*< 0.05, N = 4/group). D -Log(FDR) values for selected biological processes from the gene ontology analysis of the up- and downregulated proteins between 3 a.m. and 3 p.m.

**Supplementary Figure S8:**
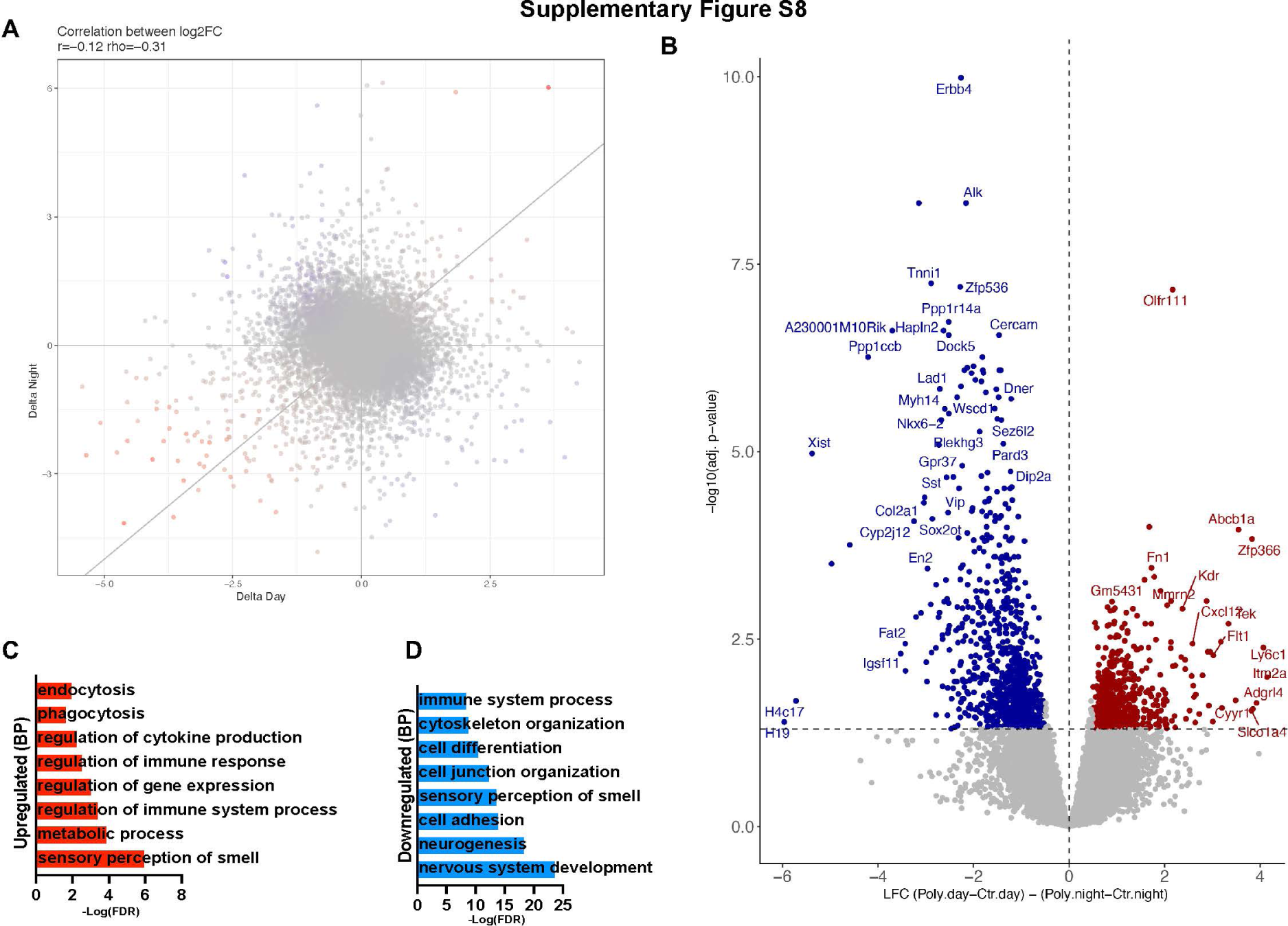
The effect of PolyI:C on the transcriptional changes between wake and sleep in adult MIA offspring. A Disco plot for the correlation between the absolute Log2-fold changes in gene expression between Poly and control animals during the active phase (x-axis) and sleep phase (y-axis). B Volcano plot showing genes significantly regulated as an effect of PolyI:C on the adult microglial transcriptional differences between the active and sleep phases (Adj. *p*< 0.05). C-D -log(FDR) values for selected biological processes associated with genes found up- (C) and downregulated (D) between the wake and sleep phase as an effect of the maternal immune activation. Images created with Biorender.com.

