## Supplementary Tables for "Microglia undergo transcriptional, translational and functional adaptations to dark and light phases in laboratory mice": Table S2.pdf

| Gene | Log2FC (1 a.m. vs. 1 p.m.) | Adj. *p* -value |
|---|---|---|
| ***Fabp3*** | **2.01** | **2E-06** |
| ***Spp1*** | **1.27** | **1.4E-03** |
| ***Gas7*** | **0.52** | **1.2E-02** |
| ***Apbb2*** | **0.52** | **1.0E-15** |
| ***Lpl*** | **0.99** | **1.6E-10** |
| ***Colec12*** | **-0.70** | **6.3E-07** |
| *Ldlr* | 0.39 | 3.5E-03 |
| *Pkm* | 0.36 | 4.7E-09 |
| *Csf1* | 0.18 | 4.7E-02 |
| ***Crybb1*** | **-0.60** | **6.1E-10** |
| *P2ry12* | -0.40 | 1.7E-04 |
| *Serpine2* | -0.32 | 1.4E-04 |
| *Ccr5* | -0.31 | 1.9E-04 |
| *P2ry13* | -0.24 | 3.7E-02 |
| *Myd88* | -0.24 | 3.2E-03 |
| *Fcrls* | -0.28 | 4.5E-03 |
| *Ctsb* | -0.21 | 3.3E-02 |
| *Cx3cr1* | -0.22 | 2.6E-02 |
| *Tmem19* | -0.17 | 4.3E-02 |
| ***Clec4e*** | **1.96** | **6.8E-03** |
| ***Clec18a*** | **1.10** | **3.3E-03** |
| ***Clec11a*** | **0.74** | **3.8E-02** |
| ***Clec1a*** | **0.68** | **4.4E-03** |
| ***Clec4b1*** | **-1.17** | **9.9E-03** |
| ***Clec5a*** | **-0.68** | **3.4E-15** |
| ***Clec4a1*** | **-0.65** | **1.8E-04** |
| *Clec4a2* | -0.35 | 1.1E-04 |
| *Clec4a3* | -0.44 | 3.9E-05 |
| In Bold: genes within the selected Log2FC cut-off of 0.5. | | |
