## Supplementary Tables for "Microglia undergo transcriptional, translational and functional adaptations to dark and light phases in laboratory mice": Table S5.pdf

| **Transcriptional Analysis** | | | | | **Proteotype Analysis** | | |
|---|---|---|---|---|---|---|---|
| **Mitochondrial Functions** | | | | | | | |
| **Gene** | **Log2FC** (1 a.m. Vs. 1 p.m.) | **Adj *p*-value** | **Log2FC** (3 a.m. Vs. 3 p.m.) | **Adj *p*-value** | **Protein** | **Log2FC** (3 a.m. Vs. 3 p.m.) | **Adj *p*-value** |
| *Ndufa10* | 0 | ns | 0.18 | ns | Q99LC3 | 0.42 | 0.037 |
| *Ndufa4* | 0.45 | 8.49E-05 | 0.13 | ns | Q62425 | -0.36 | 0.043 |
| *Ndufs1* | 0 | ns | 0 | ns | Q91VD9 | -0.36 | 0.024 |
| *Ndufb6* | 0.56 | 6.59E-06 | -0.16 | ns | Q3UIU2 | -0.73 | 0.037 |
| *Mff* | 0 | ns | 0.14 | ns | Q6PCP5 | -0.92 | 0.035 |
| **Glycolysis** | | | | | | | |
| **Gene** | **Log2FC** (1 a.m. Vs. 1 p.m.) | **Adj *p*-value** | **Log2FC** (3 a.m. Vs. 3 p.m.) | **Adj *p*-value** | **Protein** | **Log2FC** (3 a.m. Vs. 3 p.m.) | **Adj *p*-value** |
| *Slc2a1* | 2.77 | 6.97E-10 | 0.79 | **0.06** | P17809 | -1.89 | 0.034 |
| *Eno1* | 0.26 | 4.72E-04 | 0 | ns | P17182 | -1.04 | 0.030 |
| *Pgk1* | 0.63 | 5.60E-12 | 0 | ns | P09411 | -0.90 | 0.036 |
| *Pgam1* | 0.32 | 2.25E-03 | 0 | ns | Q9DBJ1 | -0.60 | 0.039 |
| *Ldha* | 0 | ns | 0 | ns | P06151 | -0.35 | 0.047 |
| *HK2* | -0.43 | 2.48E-13 | -0.44 | ns | O08528 | -0.21 | ns |
| **Lipid Metabolism** | | | | | | | |
| **Gene** | **Log2FC** (1 a.m. Vs. 1 p.m.) | **Adj *p*-value** | **Log2FC** (3 a.m. Vs. 3 p.m.) | **Adj *p*-value** | **Protein** | **Log2FC** (3 a.m. Vs. 3 p.m.) | **Adj *p*-value** |
| *Acot2* | 0.51 | 4.60E-04 | 0 | ns | Q9QYR9 | -2.20 | 0.048 |
| *Acot7* | 0.21 | ns | -0.15 | ns | Q91V12 | -1.35 | 0.037 |
| *Acad8* | 0.61 | 1.46E-19 | -0.21 | ns | Q9D7B6 | -1.00 | 0.084 |
| *Acadl* | 0.14 | ns | 0.17 | ns | P51174 | -0.58 | 0.037 |
| *Acadm* | 0 | ns | -0.15 | ns | P45952 | -0.38 | 0.037 |
| **Microglia/Macrophage activation/Integrin signalling** | | | | | | | |
| **Gene** | **Log2FC** (1 a.m. Vs. 1 p.m.) | **Adj *p*-value** | **Log2FC** (3 a.m. Vs. 3 p.m.) | **Adj *p*-value** | **Protein** | **Log2FC** (3 a.m. Vs. 3 p.m.) | **Adj *p*-value** |
| *C1qa* | -0.39 | 1.15E-04 | -0.10 | ns | P98086 | -0.39 | 0.046 |
| *C1qc* | -0.34 | 1.42E-03 | -0.14 | ns | Q02105 | -0.41 | 0.021 |
| *Itgb1* | -0.21 | 4.48E-03 | 0 | ns | P09055 | -1.02 | 0.029 |
| *Ilk* | -0.17 | 2.00E-03 | -0.18 | ns | O55222 | -0.75 | 0.044 |
| *Entpd1* | -0.76 | 7.16E-21 | -0.06 | ns | P55772 | -0.83 | 0.045 |
| *P2ry12* | -0.40 | 1.69E-04 | 0.52 | ns | Q9CPV9 | -0.27 | 0.031 |
| *Cd68* | -0.06 | ns | 0.52 | ns | P31996 | -0.22 | 0.042 |
| *Cav1* | 0.36 | ns | 1.68 | **0.08** | P49817 | -1.12 | 0.040 |
| *Grb2* | -0.25 | 2.97E-04 | 0 | ns | Q60631 | -0.40 | 0.028 |
| *Glul* | -0.33 | 3.66E-04 | -0.14 | ns | P15105 | -0.95 | 0.042 |
| *Ptgs1* | -0.14 | 4.83E-02 | -0.12 | ns | P22437 | -0.32 | 0.037 |
| *Hpgds* | -0.14 | ns | 0.24 | ns | Q9JHF7 | -0.45 | 0.047 |
| *Mapk1* | -0.11 | 4.19E-02 | 0.27 | ns | P63085 | -0.35 | 0.039 |
| *Cd81* | -0.25 | 5.05E-03 | -0.16 | ns | P35762 | -1.28 | 0.037 |
| *Hspa12a* | 0.64 | 8.64E-05 | 0.35 | ns | Q8K0U4 | -1.32 | 0.022 |
| **Motility/Phagocytosis/Cytoskeleton Polymerization** | | | | | | | |
| **Gene** | **Log2FC** (1 a.m. Vs. 1 p.m.) | **Adj *p*-value** | **Log2FC** (3 a.m. Vs. 3 p.m.) | **Adj *p*-value** | **Protein** | **Log2FC** (3 a.m. Vs. 3 p.m.) | **Adj *p*-value** |
| *Cct2* | -0.23 | 3.47E-03 | 0 | ns | P80314 | -0.44 | 0.039 |
| *Cct5* | -0.42 | 6.92E-10 | -0.12 | ns | P80316 | -0.45 | 0.043 |
| *Cct3* | -0.36 | 1.45E-09 | -0.16 | ns | P80318 | -0.46 | 0.020 |
| *Cdc42* | -0.28 | 9.91E-04 | 0.07 | ns | P60766 | -1.18 | 0.049 |
| *Dock10* | -0.46 | 1.19E-08 | -0.12 | ns | Q8BZN6 | 0.99 | 0.024 |
| *Myo1c* | 0.00 | ns | 0 | ns | Q9WTI7 | -1.06 | 0.049 |
| *Myh14* | 0.90 | 3.87E-04 | -1.84 | 4.58E-05 | Q6URW6 | -0.70 | 0.046 |
| *Myh10* | 0.28 | ns | 0.20 | ns | Q61879 | -1.68 | 0.029 |
| *Cfl2* | -0.01 | ns | 0.23 | ns | P45591 | -1.12 | 0.026 |
| *Pfn2* | 0.72 | 7.85E-05 | -0.02 | ns | Q9JJV2 | -1.32 | 0.032 |
| *Pxn* | -0.10 | 3.80E-02 | -0.11 | ns | Q8VI36 | -1.39 | 0.045 |
| *Coro2b* | 0.49 | 6.43E-03 | 0.13 | ns | Q8BH44 | -1.50 | 0.025 |
| **Extracellular Matrix Components** | | | | | | | |
| **Gene** | **Log2FC** (1 a.m. Vs. 1 p.m.) | **Adj *p*-value** | **Log2FC** (3 a.m. Vs. 3 p.m.) | **Adj *p*-value** | **Protein** | **Log2FC** (3 a.m. Vs. 3 p.m.) | **Adj *p*-value** |
| *Agrn* | 0.47 | 5.19E-03 | 0 | ns | A2ASQ1 | -1.60 | 0.036 |
| *Hspg2* | 1.10 | 5.92E-07 | 0.69 | ns | Q05793 | -1.08 | 0.029 |
| *Col4a2* | 0.06 | ns | 0 | ns | P08122 | -1.14 | 0.044 |
| *Lama2* | 0.73 | 3.03E-05 | -0.55 | ns | Q60675 | -1.28 | 0.042 |
| *Lamc1* | 0.27 | 2.22E-02 | 0.03 | ns | P02468 | -1.21 | 0.029 |
| *Nid2* | 0.58 | 7.55E-16 | 0.59 | ns | O88322 | -1.22 | 0.029 |
| **Chromatin Remodelling Complexes (NuRD and SWI/SNF)** | | | | | | | |
| **Gene** | **Log2FC** (1 a.m. Vs. 1 p.m.) | **Adj *p*-value** | **Log2FC** (3 a.m. Vs. 3 p.m.) | **Adj *p*-value** | **Protein** | **Log2FC** (3 a.m. Vs. 3 p.m.) | **Adj *p*-value** |
| Smarca5 | -0.10 | **0.079** | -0.24 | ns | Q91ZW3 | 0.75 | 0.028 |
| Smarcd1 | 0.09 | ns | 0.07 | ns | Q61466 | 0.74 | 0.029 |
| Smarcb1 | 0.07 | ns | -0.12 | ns | Q9Z0H3 | 0.65 | 0.037 |
| Arid2 | -0.27 | 3.57E-08 | 0.06 | ns | E9Q7E2 | 0.51 | 0.031 |
| Mta2 | -0.12 | 0.007 | -0.18 | ns | Q9R190 | 0.28 | 0.050 |
| **Epigenetic Readers/Chromatin Remodeling Complex Recruiters** | | | | | | | |
| **Gene** | **Log2FC** (1 a.m. Vs. 1 p.m.) | **Adj *p*-value** | **Log2FC** (3 a.m. Vs. 3 p.m.) | **Adj *p*-value** | **Protein** | **Log2FC** (3 a.m. Vs. 3 p.m.) | **Adj *p*-value** |
| Chd2 | -0.04 | ns | 0.04 | ns | E9PZM4 | 1.27 | 0.046 |
| Brd4 | -0.14 | 0.01 | 0 | ns | Q9ESU6 | 1.27 | 0.027 |
| Cbx5 | 0.36 | 4.53E-05 | 0.09 | ns | Q61686 | 0.95 | 0.020 |
| Cbx8 | -0.02 | ns | 0 | ns | Q9QXV1 | 0.57 | 0.029 |
| Mbd1 | -0.04 | ns | -0.09 | ns | Q9Z2E2 | 0.88 | 0.034 |
| **Epigenetic Writers (DNA-methyltransferase/Histone demethylase)** | | | | | | | |
| **Gene** | **Log2FC** (1 a.m. Vs. 1 p.m.) | **Adj *p*-value** | **Log2FC** (3 a.m. Vs. 3 p.m.) | **Adj *p*-value** | **Protein** | **Log2FC** (3 a.m. Vs. 3 p.m.) | **Adj *p*-value** |
| Mecp2 | -0.03 | ns | 0.09 | ns | Q9Z2D6 | 0.75 | 0.026 |
| Kdm5b | -0.06 | ns | -0.17 | ns | Q80Y84 | 0.80 | 0.037 |
