## Supplementary Tables for "Microglia undergo transcriptional, translational and functional adaptations to dark and light phases in laboratory mice": Table S6.pdf

| | Protein | Gene | Baseline Night vs. Day | LPS Day | | LPS Night | |
|---|---|---|---|---|---|---|---|
| | | | Log2FC Protein | Log2FC RNA | Log2FC Protein | Log2FC RNA | Log2FC Protein |
| SWI/SNF Remodeling Complex | Q3TKT4 | Smarca4 | 0.28 | -0.16 | **0.78** | -0.1 | 0.04 |
| | Q6PGB8 | Smarcal1 | *1.01* | -0.12 | **1.58** | 0.04 | 0.57 |
| | Q91ZW3 | Smarca5 | **0.75** | **-0.79** | **1.00** | **-0.77** | *0.46* |
| | Q61466 | Smarcd1 | **0.74** | 0.16 | **1.13** | **0.38** | 0.26 |
| | Q6PGB8 | Smarca1 | *1.01* | 0.42 | **1.58** | 0.41 | 0.57 |
| | E9Q7E2 | Arid2 | **0.51** | 0.08 | **0.90** | -0.12 | -0.07 |
| | A2BH40 | Arid1a | 0.40 | 0.12 | **0.71** | *-0.41* | 0.06 |
| | E9Q4N7 | Arid1b | 0.30 | 0.18 | **0.97** | **0.35** | 0.13 |
| NuRD Remodeling Complex | P97868 | Rbbp6 | **0.70** | **-0.63** | **1.33** | **-0.44** | 0.31 |
| | Q8BX09 | Rbbp5 | 0.34 | **-0.25** | **0.59** | **-0.36** | 0.19 |
| | O88851 | Rbbp9 | 0.10 | **0.57** | -0.59 | 0.40 | -0.45 |
| | Q9R190 | Mta2 | **0.28** | *-0.28* | **0.54** | **-0.45** | **0.29** |
| | Q924K8 | Mta3 | **0.44** | **0.34** | **0.88** | **0.61** | **0.51** |
| Epigenetic Mark Readers | Q6PDQ2 | Chd4 | 0.85 | *-0.31* | **1.33** | *-0.33* | 0.54 |
| | A2A8L1 | Chd5 | *0.67* | 0.39 | **1.00** | **0.89** | 0.35 |
| | E9PZM4 | Chd2 | **1.27** | **-0.31** | **1.81** | -0.15 | 0.74 |
| | Q7JJ13 | Brd2 | 0.62 | -0.25 | **0.93** | **-0.45** | 0.42 |
| | Q9ESU6 | Brd4 | **1.27** | -0.18 | **2.12** | -0.24 | 0.50 |
| | Q8K2F0 | Brd3 | *0.40* | **0.43** | **0.97** | **0.24** | *0.41* |
| | Q9QXV1 | Cbx5 | **0.57** | **-0.92** | **1.49** | **-0.84** | -0.18 |
| | Q9Z2D6 | Mecp2 | **0.75** | *-0.25* | **1.30** | 0.09 | *0.36* |
| | Q9Z2E2 | Mbd1 | **0.88** | -0.07 | **0.93** | -0.09 | *-0.56* |
| Epigenetic Writers (Histone/DNA Modifications) | O88508 | Dnmt3a | 0.49 | **0.69** | **1.11** | **0.60** | 0.14 |
| | Q3UXZ9 | Kdm5a | *0.41* | -0.09 | **1.11** | 0.07 | 0.47 |
| | Q80Y84 | Kdm5b | **0.80** | **0.65** | **1.34** | **-0.17** | 0.21 |
| | O09106 | Hdac1 | 0.19 | -0.25 | 0.04 | *-0.36* | **-0.65** |
| | P70288 | Hdac2 | 0.39 | **-0.40** | **0.74** | **-0.57** | -0.12 |

**Bold**: Adj p-value <0.05, *Italic*: Trend (Adj p-value: 0.055-0.09),Normal: non-significant
